# Glycoengineering of the hepatitis C virus E2 glycoprotein leads to improved biochemical properties and enhanced immunogenicity

**DOI:** 10.1101/2025.04.02.646860

**Authors:** Liudmila Kulakova, Kiki H. Li, Austin W. T. Chiang, Michael P. Schwoerer, Sanne Schoffelen, Khadija Elkholy, Kinlin L. Chao, Salman Shahid, Bhoj Kumar, Nathan B. Murray, Stephanie Archer-Hartmann, Parastoo Azadi, Bjørn G. Voldborg, Alexander Marin, Roy A. Mariuzza, Alexander K. Andrianov, Alexander Ploss, Nathan E. Lewis, Eric A. Toth, Thomas R. Fuerst

**Author notes:** Correspondence to (T.R.F.).

## Abstract

An effective vaccine against hepatitis C virus (HCV) must elicit the production of broadly neutralizing antibodies (bnAbs) reproducibly against the E1E2 glycoprotein complex. Little is known about how glycan content affects this process. Ideally, glycans would maximize epitope exposure without compromising antigen stability or exposing new epitopes. However, typical recombinant vaccines contain considerable heterogeneity in glycan content, which can affect the antibody response and neutralization potency. Here we employed glycoengineered Chinese hamster ovary (geCHO) cell lines that impart nearly homogeneous glycosylation as a means to test how specific glycan features influence antigenicity and immunogenicity for the secreted HCV E2 ectodomain (sE2). Specific geCHO antigens exhibited a modest but reproducible increase in affinity for some mAbs relative to CHO- and HEK293-produced sE2. Surprisingly, one geCHO sE2 antigen failed to bind the CD81 receptor, indicating the potential for significant glycan effects on biochemical properties. We immunized mice with the four antigens and found the total antibody response to be the same for all groups. However, sera from one geCHO group exhibited a 7-fold improvement in neutralization against the homologous HCV pseudovirus and had the most mice whose sera exhibited neutralization activity against genotypes 1b, 2a, 2b, and 3. Further analysis identified beneficial and deleterious glycan features, and the glycan that correlated the most with decreased potency was relatively small. However, size was not the sole determinant of glycan-driven effects on the antibody response. In summary, glycan content impacts biochemical properties of antigens to varying degrees and such effects can influence immune response quality and uniformity.

## INTRODUCTION

HCV is a major cause of severe liver diseases and cancer. More than 50 million individuals are chronically infected, with an annual increase of 1 million new infections ^1^. HCV is responsible for more deaths in the USA than all other infectious diseases combined, ^2^ with an estimated 242,000 deaths attributable to HCV in 2022. HCV infection progresses to chronic illness in nearly 75% of cases, increasing the risk for development of cirrhosis or hepatocellular carcinoma. The World Health Organization recently announced a global hepatitis strategy to reduce new infections of hepatitis viruses by 90% and associated deaths by 65% by 2030 ^1^. As one major component of the global hepatitis strategy, direct-acting antivirals (DAAs), which cure existing HCV infections, ^3, 4^ will significantly aid in reaching the WHO target. However, DAA-treated individuals can become re-infected which decreases effectiveness in high-risk groups. Moreover, most HCV-infected individuals are asymptomatic until liver damage is extensive enough to present as liver disease ^1^. Therefore, an effective prophylactic HCV vaccine is essential to the WHO global strategy to reduce the burden of viral hepatitis disease ^5^.

HCV is an RNA virus that rapidly accumulates mutations, leading to considerable genetic diversity and making vaccine development challenging. However, spontaneous clearance rates and the presence of immune memory among individuals who clear their first HCV infection ^6–9^ suggest that an HCV vaccine is feasible ^5, 10–12^. HCV contains two glycoproteins, E1 and E2, that comprise the envelope glycoprotein complex. E1 and E2 are processed in the endoplasmic reticulum (ER) of liver cells by peptidases and cellular glycosylation machinery to produce the mature E1E2 complex. E1 and E2 are type I transmembrane proteins with a highly glycosylated N-terminal ectodomain and a C-terminal hydrophobic anchor that forms a membrane-anchored complex, mbE1E2, on the surface of the virion. The mbE1E2 heterodimer interacts with host cell receptors to mediate viral entry into the cell, making E1E2 the prime candidate for an HCV vaccine. Both a soluble version of the E2 protein (sE2) lacking the transmembrane domain ^13 14–20^ and the E1E2 complex in both secreted and membrane-bound forms have been the subject of vaccine studies. Immunological assessment in guinea pigs ^21^ and chimpanzees suggest that an E1E2-based vaccine is superior to E2 administered alone ^22–24^, but the E2 ectodomain remains a useful model system due to its ease of production and the presence of the receptor-binding interface, which contains epitopes that give rise to neutralizing antibodies (nAbs) ^25–27^. In particular, a recent study using plasma deconvolution to assess nAb populations in patients with persistent HCV infection versus patients that cleared the infection showed that clearance is associated with the presence of multiple bnAb types in the polyclonal sera ^28^. Moreover, the bnAb combinations most highly associated with clearance included an E2-specific bnAb (HEPC74, AR3A, or HC84.26) in combination with the E1E2-specific bnAb AR4A.

The E1 and E2 proteins are decorated with extensive host post-translational modifications, including a prominent glycan shield^29, 30^ that hides antigenic epitopes from nAbs^31, 32^ (**Figure 1**). E1 and E2 have up to 5 and 11 glycosylation sites, respectively. These are mostly conserved across HCV lineages^33–35^, including glycans flanking the E2 antigenic face^36^. The glycans reduce the sensitivity of cell culture-derived HCV (HCVcc) to antibody neutralization^37, 38^. In addition, some glycans are important for viral entry^37, 38^ and E1E2 complex assembly^35^. N-linked glycans influence both local and global structure in subtle ways, based on a survey of modified versus unmodified structure pairs in the PDB^39^. Moreover, glycans can influence antibody affinity. Extensive studies of the anti-cancer target MUC1 showed with synthetic peptides and glycopeptides harboring different glycans that differences in glycan structure can have varying effects on antibody affinity, either increasing, decreasing, or having no effect depending on the glycoform^40^. Thus, similar precise tuning of glycan content could influence the antigenicity of HCV envelope glycoprotein vaccine candidates. Toward that end, previous work by our group and others using sE2 expressed in HEK 293 versus Sf9 cells^17, 19^ showed similar effects, but were less systematic and lacked the data to draw specific conclusions, due to the inherent glycoform heterogeneity in antigens produced in those systems. By contrast, glycan removal can create neoepitopes that elicit antibodies that do not recognize the native virus, as observed for HIV-1 env^41^ as well as potential destabilization of the protein structure. Glycosylation can impact antigen uptake, proteolytic processing, and MHC presentation, all of which determine the cellular immune response^42^. Indeed, enzymatically trimming all glycans on influenza hemagglutinin and SARS-CoV-2 Spike protein to a single GlcNAc resulted in more potent neutralization and enhanced survival in a challenge model^43, 44^, and stronger humoral and cell-mediated immune responses^43, 44^. The modified spike protein also afforded better protection against SARS CoV-2 challenge in a humanized mouse model^43^. Similarly, trimming glycans on HIV env resulted in more frequent vaccine responders in a mouse model^45^. These data suggest that glycan optimization could be important to vaccine design and optimization.

**Figure 1.**
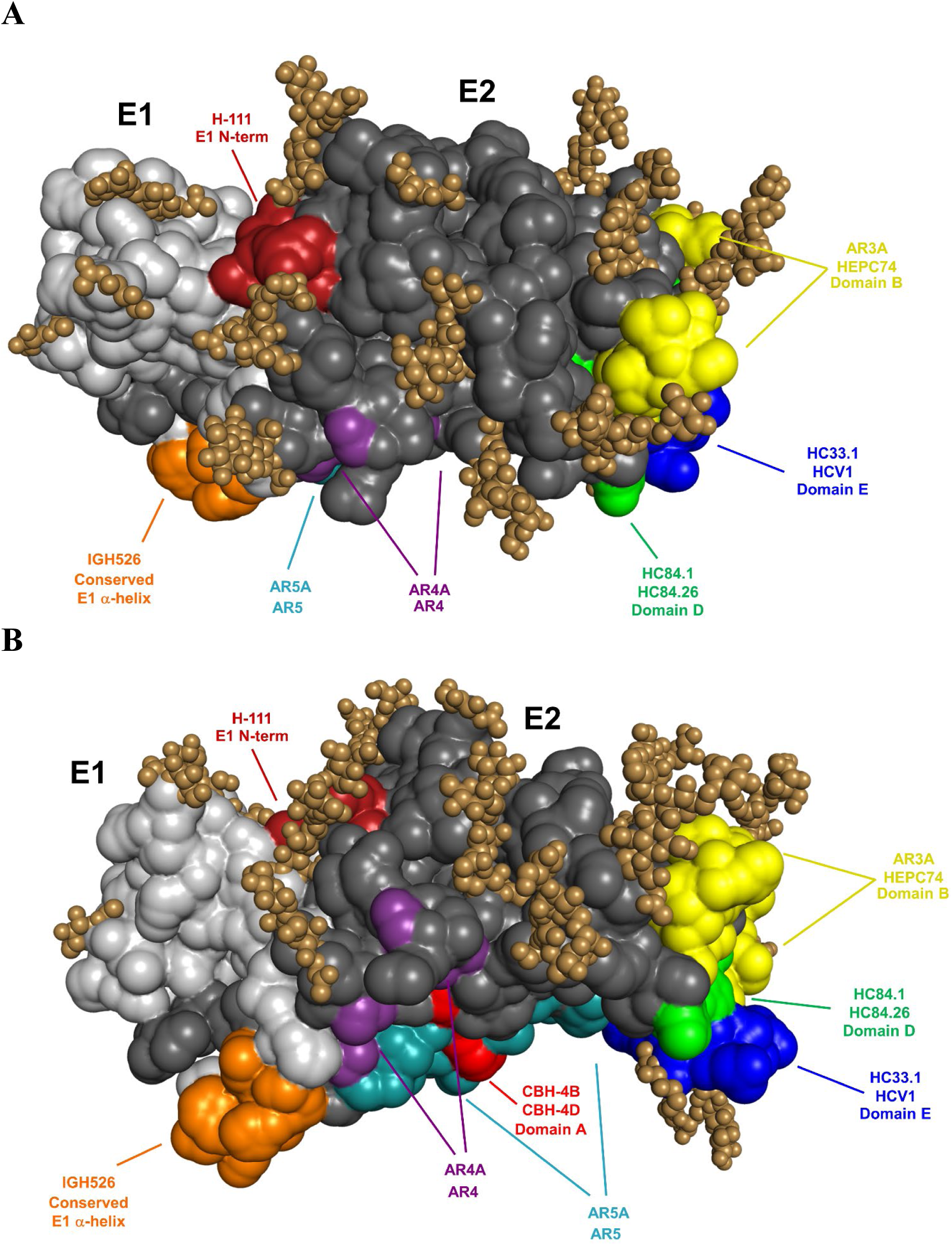
**A.** Surface depiction of the E1E2 complex with antigenic domains highlighted and representative antibodies that bind to those domains identified. E2 antigenic domain A is colored red, domain B is colored yellow, domain D is colored green, domain E is colored blue, AR4 is colored purple, and AR5 is colored cyan. The E1 antigenic domain consisting of the N-terminus is colored brick red, and the conserved helix is colored orange. Glycans surrounding these antigenic domains are depicted as sand-colored spheres. **B**. A view of the complex rotated 60 degrees relative to **A** around an axis parallel to the plane of the screen.

Mammalian cells have many glycosyltransferases synthesizing glycan structures on proteins, including a few dozen enzymes making N-linked glycans on E1E2 heterodimers. Chinese hamster ovary (CHO) cells express most glycosyltransferases used in mammalian N-linked glycosylation^46, 47^, including enzymes for complex/hybrid N-glycans. However, the enzymes synthesize the glycans through complex pathways that are difficult to control. Thus, for a glycoprotein, often multiple glycoforms are made, with differences in glycan structures decorating the protein depending on expression host, as reported for sE2 ^17, 19^. Some of these differences are substantial, even between glycans of similar abundance at the same glycosylation site on the antigen^19^. Unfortunately, this heterogeneity leads to the presence of suboptimal glycans which dilute vaccine potency. Thus, an ideal vaccine would contain only the most potent glycoforms and be produced in a cell line engineered to append the favored glycan(s) to the candidate vaccine. To develop a biomanufacturing system that allows precision control of mammalian glycoforms, we deeply characterized glycosylation in CHO cells. First, we sequenced the CHO-K1 and Chinese hamster genomes and quantified glycosyltransferase mRNA expression^46, 48–50^. Further, we and others linked each glycosyltransferase to their functions by knocking out each gene individually and in combination and then quantifying the glycome and mRNA, and using specialized systems biology modeling techniques to confirm their role in glycosylation and identify indirect effects^47, 51–53^. We used these data to build 30 glycoengineered CHO (geCHO) cell lines, wherein we knocked out/in 17 genes, alone or in combination, from CHO-S cells^54^.

Here we report the production, biochemical characterization, and in vivo immunogenicity of specific glycoforms HCV sE2, a main target of the antibody response to HCV. We tested sE2 produced in wild-type CHO, human embryonic kidney (HEK) cells, and our geCHO cells. Following an initial screen of twelve geCHO sE2 glycoforms, we chose two with enhanced binding to bnAbs against HCV. These glycoforms showed improved homogeneity when compared to the CHO- and HEK-derived antigens. Remarkably, one glycoform eliminated binding to the HCV entry receptor CD81, demonstrating an unexpected ability of glycan content to regulate envelope glycoprotein function. Assessment of sera from mice immunized with these antigens showed comparable antibody titers and levels of domain-specific antibodies to conserved, continuous and discontinuous epitopes. Remarkably, the sera from the geCHO groups showed a tighter clustering of IC_50_ values compared to the CHO and HEK groups. Moreover, one glycoform, geCHO.sE2.1, elicited a more potent neutralizing response than the other three groups and exhibited a greater frequency of neutralizing activity against heterologous HCV pseudoviruses. Further analysis of glycans on each glycoform elucidated glycan features that contributed both positively and negatively to the immune response. Thus, different glycoforms have distinct effects on both biochemical properties and the immune responses. Furthermore, the size of the glycoform is not the sole determinant of these effects. Finally, this underscores the benefit of incorporating glycan optimization into vaccine design pipelines.

## RESULTS

### Initial screen of HCV E2 glycovariants

To prioritize potential glycoform candidates for biochemical and immunological analysis, we first produced small batches of sE2 in twelve different geCHO cell lines (**Figure S1**), plus CHO-S and HEK 293, each of which were subsequently glycoprofiled (see **Figures S3** and **S6**). The glycoforms geCHO.sE2.3, geCHO.sE2.9, geCHO.sE2.11, and geCHO.sE2.12 were eliminated due to relatively low expression levels (**Figure S1**). Binding of nAbs to the geCHO antigens was analyzed by low-volume ELISA (due to the limited quantities of some of the sE2 glycoforms; see **Figure S1**) and benchmarked against the sE2 produced HEK 293, which is well-characterized (**Table 1**) ^15, 18, 19, 55^ and has been used by our group in several studies ^15, 18, 19, 55^. This initial screen identified five glycoforms that exhibited enhanced binding to the bnAbs AR3A, which recognizes antigenic domain B ^56^, and HC84.26, which recognizes antigenic domain D ^57^. These five glycoforms were geCHO.sE2.1, geCHO.sE2.2, geCHO.sE2.4, geCHO.sE2.6, and geCHO.sE2.7. The group of five glycoforms comprise two structurally related sets, and two candidate glycoforms from the set were chosen as follows. First, geCHO.sE2.1 was chosen in favor of geCHO.sE2.2 and geCHO.sE2.4 due to a combination of expression levels and antibody binding (**Figure S1** and **Table 1**). The geCHO.sE2.1 antigen exhibited higher expression levels than geCHO.sE2.2 and superior binding to both AR3A and HC84.26 than geCHO.sE2.4. For the geCHO.sE2.6/geCHO.sE2.7 pair, binding to AR3A and HC84.26 was indistinguishable, but geCHO.sE2.7 exhibited substantially higher expression levels and was thus chosen. The dominant glycan in the geCHO.sE2.7 antigen is also the dominant glycoform appended to proteins secreted by the liver ^58^, which also factored into its choice as the second candidate glycoform. To determine whether the bnAb binding differences were due to the imparted glycans, we expressed and purified geCHO.sE2.1 and geCHO.sE2.7 at a larger scale for deglycosylation and re-analysis. The differences in antibody binding are present in the purified, glycosylated geCHO antigens relative to HEK sE2, but is no longer evident upon deglycosylation of the antigens (**Table S1 and Figure S2**).

**Table 1.**
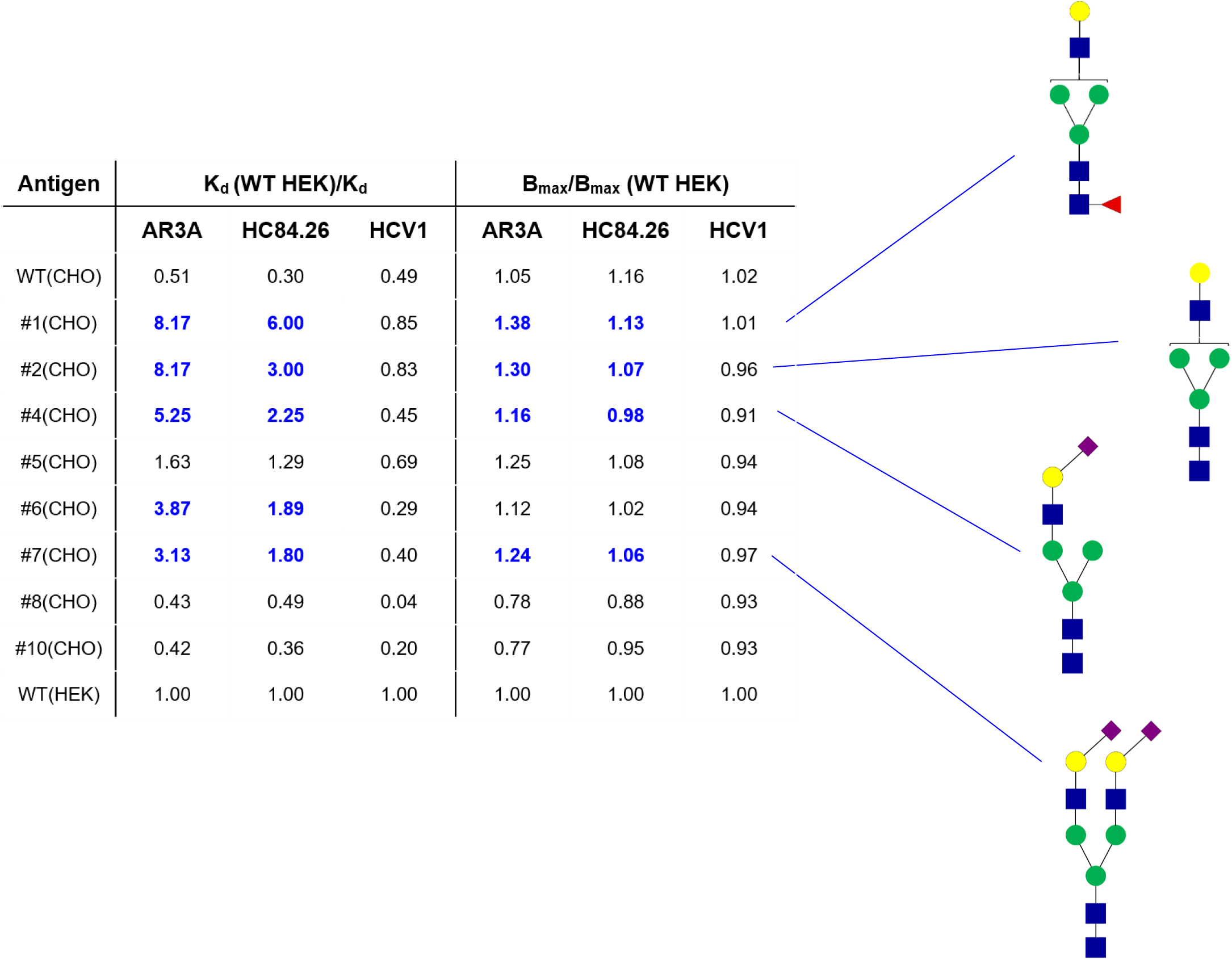
Analysis of sE2 glycoform antigenicity.

### Glycan profiles of E2 antigens

We subjected the two test antigens (geCHO.sE2.1 and geCHO.sE2.7) and control antigens produced in CHO-S and HEK 293 to N-glycomics analysis (**Tables 2-5** and **Figures S3-6**). In general, as expected the test antigens contained small glycans. In particular, for the geCHO.sE2.1 antigen, greater than 90% of the glycans present were 2605 Da or smaller, with the predominant mono-antennary non-sialylated glycan with a mass of 1794 Da comprising 36.2% of the total measured glycan population (**Table 3** and **Figure S4**). For the geCHO.sE2.7 antigen, greater than 80% of the glycan population was of a mass 2792 Da or smaller (**Table 4** and **Figure S5**), with the most abundant glycan being a biantennary sialylated glycan with a mass of 2792 Da, which comprises 23% of the overall glycan population. Larger glycans for the two test antigens are essentially absent. For geCHO.sE2.1, only 0.4% of the total glycan population is tri-or tetraantennary and for geCHO.sE2.7 this percentage is 4.6%. By contrast, the antigens derived from the typically-used CHO and HEK expression systems contain a significant amount of large glycans, with 34% of the total glycan population of the CHO-S antigen contains tri- or tetraantennary glycans (**Table 2**) as does ∼11-20% of the HEK-derived sE2. Thus, the glycan population for both test antigens is markedly skewed towards smaller glycans relative to the control antigens.

**Table 2.**
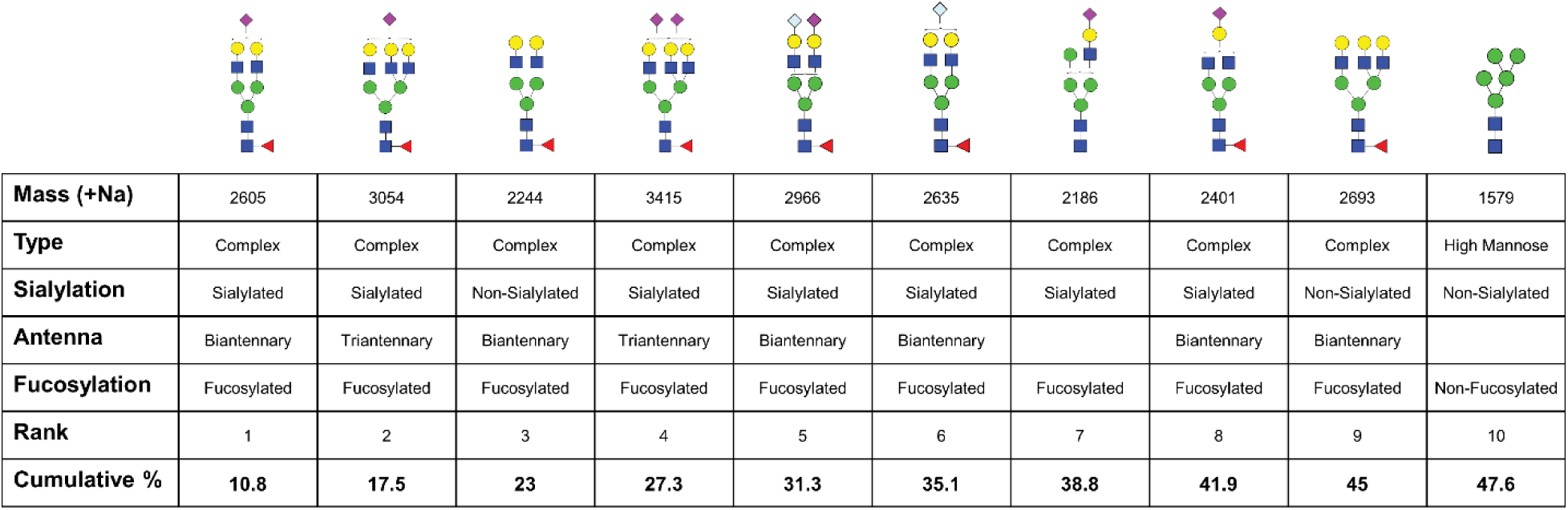
Top ten glycoforms of CHO sE2.

|  |  |  |  |  |  |  |  |  |  |  |
| --- | --- | --- | --- | --- | --- | --- | --- | --- | --- | --- |
| Mass (+Na) | 2605 | 3054 | 2244 | 3415 | 2966 | 2635 | 2186 | 2401 | 2693 | 1579 |
| Type | Complex | Complex | Complex | Complex | Complex | Complex | Complex | Complex | Complex | High Mannose |
| Sialylation | Sialylated | Sialylated | Non-Sialylated | Sialylated | Sialylated | Sialylated | Sialylated | Sialylated | Non-Sialylated | Non-Sialylated |
| Antenna | Biantennary | Triantennary | Biantennary | Triantennary | Biantennary | Biantennary |  | Biantennary | Biantennary |  |
| Fucosylation | Fucosylated | Fucosylated | Fucosylated | Fucosylated | Fucosylated | Fucosylated | Fucosylated | Fucosylated | Fucosylated | Non-Fucosylated |
| Rank | 1 | 2 | 3 | 4 | 5 | 6 | 7 | 8 | 9 | 10 |
| Cumulative % | 10.8 | 17.5 | 23 | 27.3 | 31.3 | 35.1 | 38.8 | 41.9 | 45 | 47.6 |

**Table 3.**
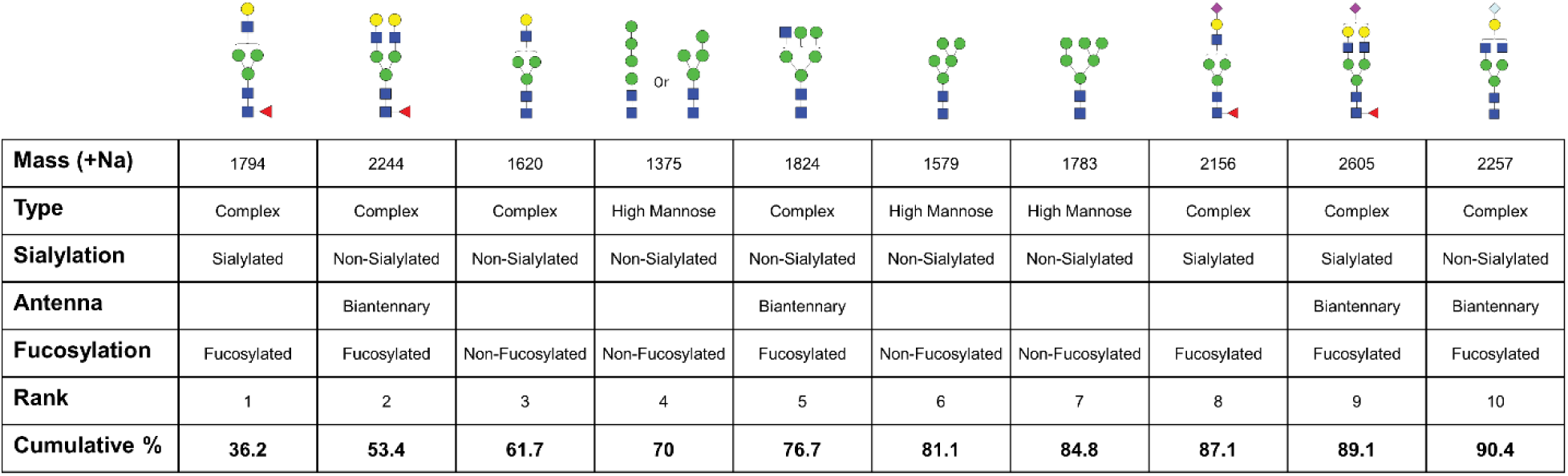
Top ten glycoforms of geCHO.sE2.1.

|  |  |  |  |  |  |  |  |  |  |  |
| --- | --- | --- | --- | --- | --- | --- | --- | --- | --- | --- |
| Mass (+Na) | 1794 | 2244 | 1620 | 1375 | 1824 | 1579 | 1783 | 2156 | 2605 | 2257 |
| Type | Complex | Complex | Complex | High Mannose | Complex | High Mannose | High Mannose | Complex | Complex | Complex |
| Sialylation | Sialylated | Non-Sialylated | Non-Sialylated | Non-Sialylated | Non-Sialylated | Non-Sialylated | Non-Sialylated | Sialylated | Sialylated | Non-Sialylated |
| Antenna |  | Biantennary |  |  | Biantennary |  |  |  | Biantennary | Biantennary |
| Fucosylation | Fucosylated | Fucosylated | Non-Fucosylated | Non-Fucosylated | Fucosylated | Non-Fucosylated | Non-Fucosylated | Fucosylated | Fucosylated | Fucosylated |
| Rank | 1 | 2 | 3 | 4 | 5 | 6 | 7 | 8 | 9 | 10 |
| Cumulative % | 36.2 | 53.4 | 61.7 | 70 | 76.7 | 81.1 | 84.8 | 87.1 | 89.1 | 90.4 |

**Table 4.** Top ten glycoforms of geCHO.sE2.7.

|  |  |  |  |  |  |  |  |  |  |  |
| --- | --- | --- | --- | --- | --- | --- | --- | --- | --- | --- |
| Mass (+Na) | 2792 | 2547 | 2227 | 2431 | 1661 | 1981 | 2186 | 2192 | 2216 | 2472 |
| Type | Complex | Complex | Complex | Complex | Complex | Complex | Complex | High Mannose | Complex | Complex |
| Sialylation | Sialylated | Sialylated | Sialylated | Sialylated | Non-Sialylated | Sialylated | Sialylated | Non-Sialylated | Sialylated | Non-Sialylated |
| Antenna | Biantennary |  | Biantennary | Biantennary | Biantennary |  |  |  |  | Triantennary |
| Fucosylation | Non-Fucosylated | Non-Fucosylated | Non-Fucosylated | Non-Fucosylated | Non-Fucosylated | Non-Fucosylated | Fucosylated | Non-Fucosylated | Non-Fucosylated | Non-Fucosylated |
| Rank | 1 | 2 | 3 | 4 | 5 | 6 | 7 | 8 | 9 | 10 |
| Cumulative % | 23 | 37.1 | 46.8 | 55 | 61.4 | 66.5 | 71.6 | 74.9 | 77.8 | 80.6 |

**Table 5.** Top ten glycoforms of HEK sE2.

|  |  |  |  |  |  |  |  |  |  |  |
| --- | --- | --- | --- | --- | --- | --- | --- | --- | --- | --- |
| Mass (+Na) | 1579 | 1335 | 2081 | 1417 | 1836 | 1662 | 1784 | 1988 | 1539 | 1621 |
| Type | High Mannose | High Mannose | Complex | Complex | Complex | Complex | High Mannose | High Mannose | High Mannose | Complex |
| Sialylation | Non-Sialylated | Non-Sialylated | Non-Sialylated | Non-Sialylated | Non-Sialylated | Non-Sialylated | Non-Sialylated | Non-Sialylated | Non-Sialylated | Non-Sialylated |
| Antenna |  |  | Triantennary |  | Biantennary | Biantennary |  |  |  |  |
| Fucosylation | Non-Fucosylated | Non-Fucosylated | Fucosylated | Non-Fucosylated | Fucosylated | Non-Fucosylated | Non-Fucosylated | Non-Fucosylated | Non-Fucosylated | Non-Fucosylated |
| Rank | 1 | 2 | 3 | 4 | 5 | 6 | 7 | 8 | 9 | 10 |
| Cumulative % | 12.1 | 22.5 | 31.3 | 38.3 | 44.0 | 49.1 | 52.6 | 55.8 | 58.5 | 61.0 |

### Biochemical analysis of E2 antigens

We purified both the test antigens and control antigens using immobilized metal affinity chromatography (IMAC) followed by size exclusion chromatography (SEC) as described previously ^15, 18, 19^. Each antigen produced using this regimen was highly purified and migrated as a single band at a molecular weight of 70-85 kDa for CHO and HEK sE2 and 60-70 kDa for geCHO.sE2.1 and geCHO.sE2.7 (**Figure 2**). The band spread for geCHO.sE2.1 and geCHO.sE2.7 in both SDS-PAGE and reducing western is narrower, most likely due to the presence of more uniform, smaller glycoforms, then that observed for the CHO and HEK antigens. In addition, we analyzed the behavior of the antigens by non-reducing western blot and observed that the antigens migrate as two distinct species. In the geCHO.sE2.1, geCHO.sE2.7, and HEK sE2 preparations, the distribution appears to be roughly evenly split, whereas for the CHO sE2 the slower-migrating species is more populated than the faster-migrating one. It is possible that this band reflects a mix of species with large glycans and some E2 multimers which have been observed previously for HEK sE2 and sE2 produced in Sf9 cells ^19^.

**Figure 2.**
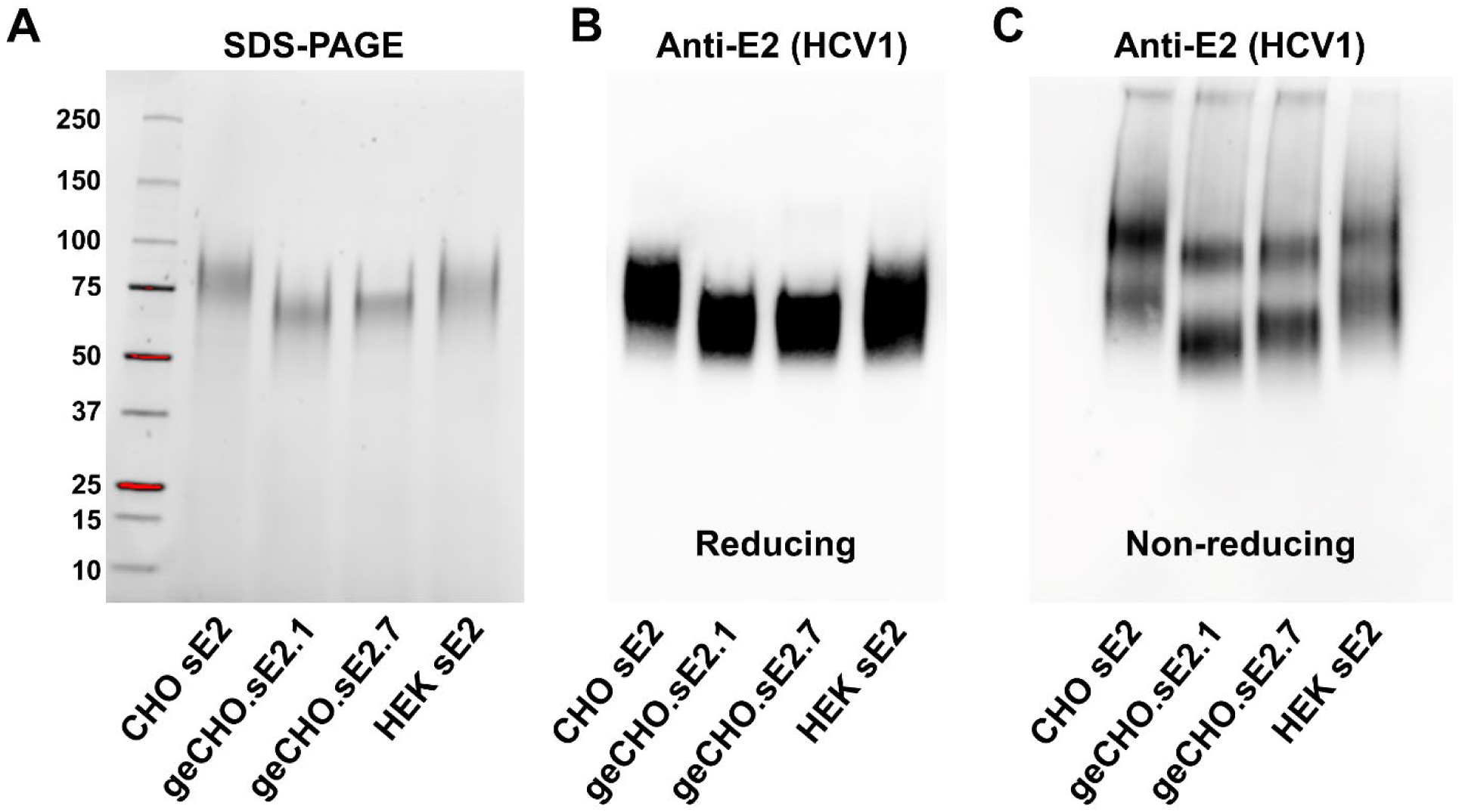
Analysis of the four sE2 glycoforms. (A) SDS-PAGE analysis of CHO sE2, geCHO.sE2.1, geCHO.sE.7, and HEK sE2 under reducing conditions. (B) Western blot detection of the purified proteins under reducing conditions. (C) Western blot detection of the purified proteins under non-reducing conditions. The anti-E2 antibody HCV1 antibody was used at a concentration of 5 μg/mL for the Western blots.

The test and control antigens were also characterized using analytical ultracentrifugation (AUC), which can separate a mixture of protein populations more precisely than SEC ^59^. A comparison of AUC results shows that all four antigens exist in solution as one major species with an additional species that we interpret as a multimeric sE2 form that is most likely dimeric (**Figure 3**). The sedimentation coefficient (S) values for the predominant peak ranges from 3.6 for geCHO.sE2.1 to 3.9 for CHO sE2, with the size difference consistent with the observed difference in glycan size in the population of glycoforms. Based on the sharpness of the peak at 3.6 S and the presence of a minor distinct peak at 5.4 S, geCHO.sE2.1 looks the most uniform. The geCHO.sE2.7 and HEK sE2 also migrate as two species, a major one (3.7 S for geCHO.sE2.7 and 3.8 S for HEK sE2) and a minor one (5.7 S for geCHO.sE2.7 and 6.4 S for HEK sE2). However, the major peaks are broader and not a separated from the minor peak as geCHO.sE2.1. The CHO sE2 sample has similar major and minor species at 3.9 S and 5.8 S respectively. The minor species for CHO sE2 is the most prominent of the four and CHO sE2 contains additional minor species at 1.8 and 8.1 S and thus appears to be the least uniform of the four antigens.

**Figure 3.**
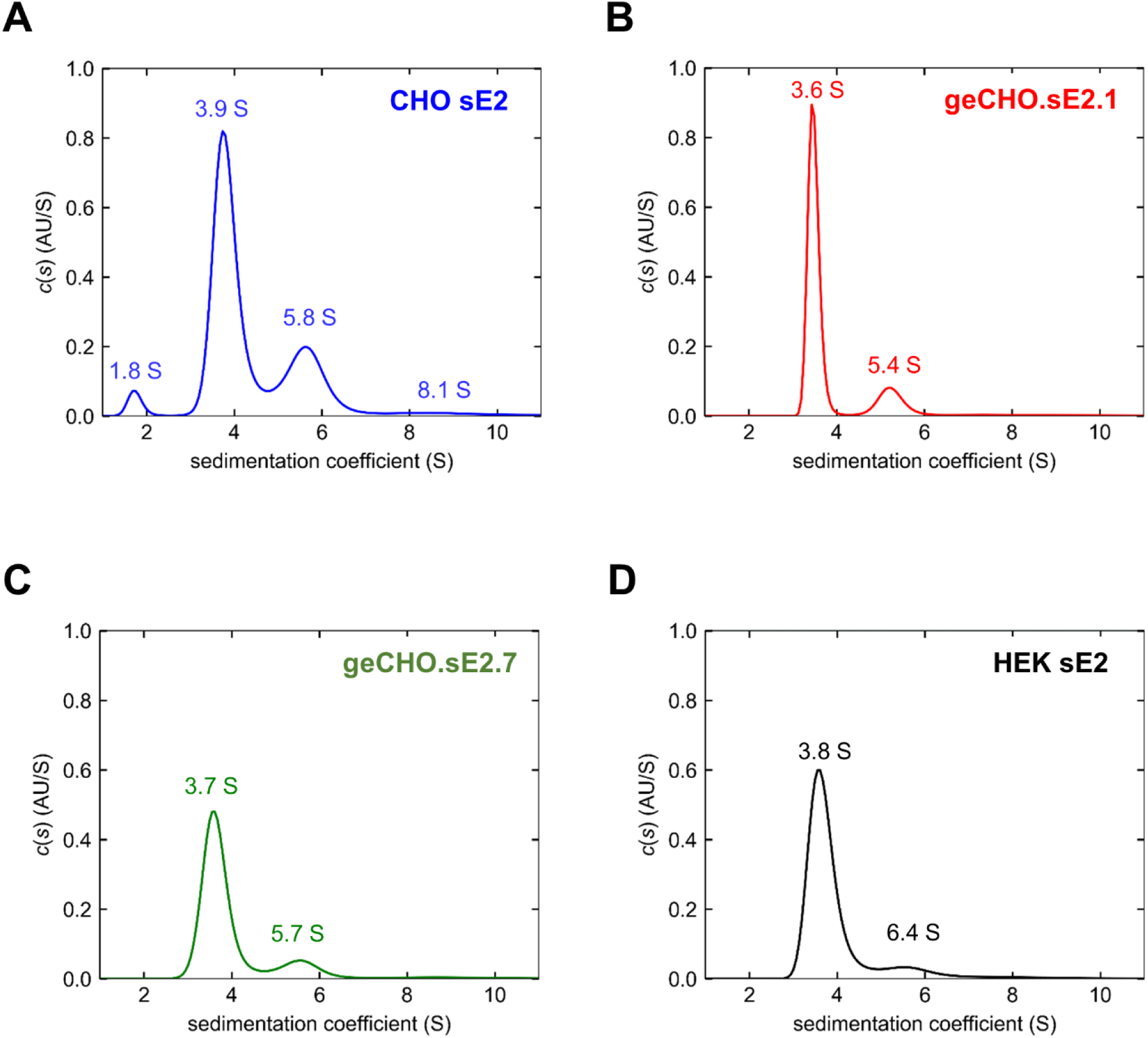
Analytical characterization of the size and heterogeneity of the four sE2 glycoforms. AUC profiles of (A) CHO sE2, (B) geCHO.sE2.1, (C) geCHO.sE2.7, and (D) HEK sE2. Shown are the distribution of Lamm equation solutions c(s) for the two proteins (solid lines). Calculated sedimentation coefficients for the peaks are labeled. All four antigens have two major species, one sedimenting at between 3.6 and 3.9 S, and on sedimenting between 5.4 and 6.4 S. The CHO sE2 sample contained two additional low-abundance species, one sedimenting at 1.8 S and the other at 8.1 S.

### Receptor binding by E2 antigens

To assess any potential effects of glycan content on binding to the primary receptor for HCV entry, CD81, we assessed binding affinity of each antigen for the CD81-LEL by ELISA. As can be seen in **Figure 4**, binding of the CHO sE2, geCHO.sE2.1, and HEK sE2 antigens to the CD81-LEL is roughly equivalent, with dissociation constants in the range of 60-80 nM. However, geCHO.sE2.7 binds the CD81-LEL poorly. The binding reaction only reaches approximately 0.4 OD at 450 nm at the highest concentration of CD81-LEL-Fc used (2.8 μM). Since the antigen binds well to all of the antibodies that surround the CD81 binding site and is well-behaved in solution, the most likely explanation is that the glycan content is in some manner interfering with CD81 binding. This surprising result highlights the potential impact of glycans on biochemical properties and underscores the need for a thorough biochemical examination for vaccine candidates in general.

**Figure 4.**
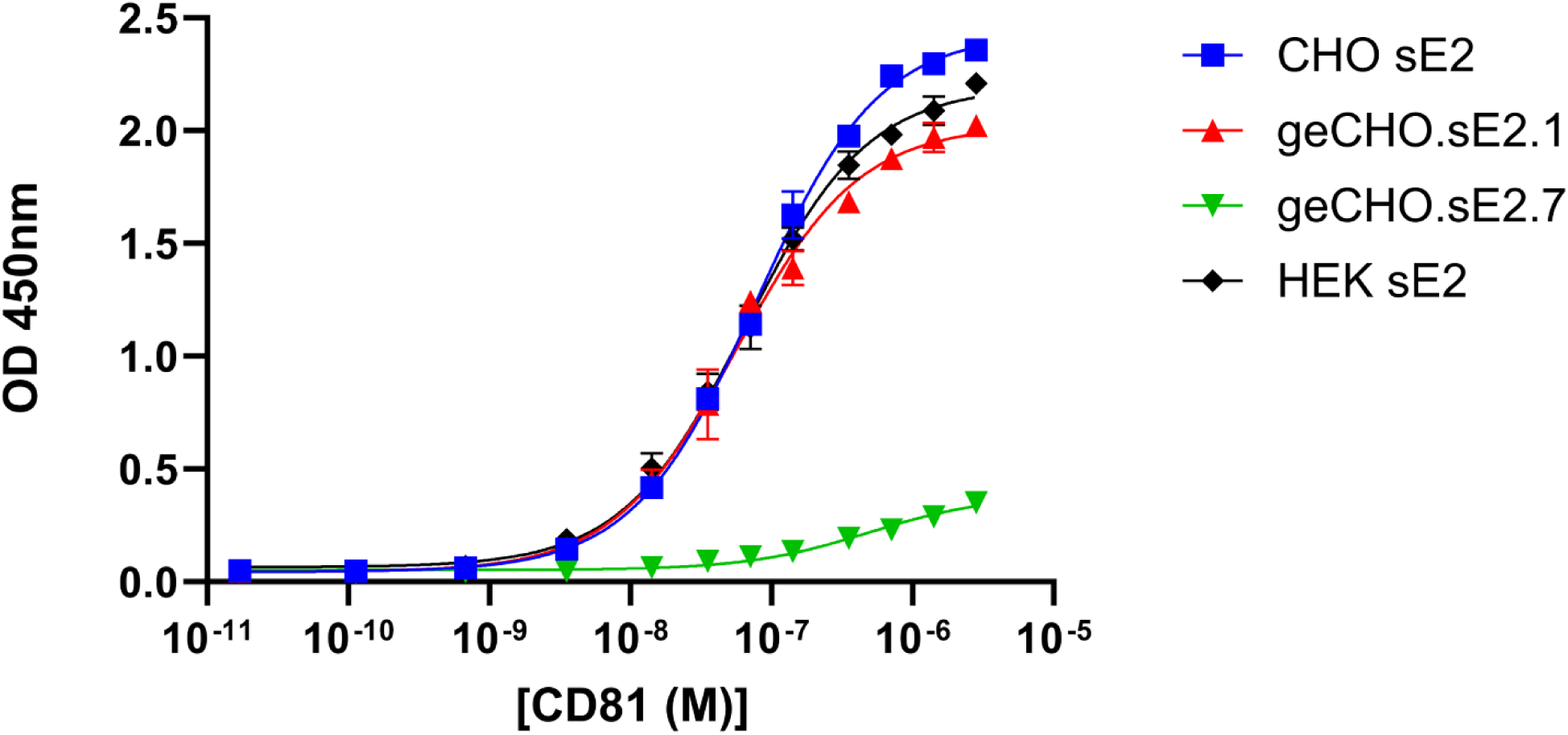
Analysis of receptor binding by ELISA. Each datum represents the mean of two duplicate experiments for each antigen. The curve overlaid on the data (*solid line*) represents the best fit of the data to a binary binding model which estimates the Kd of the interaction. Where error bars are not visible, this indicates that the error was too small relative to the symbol used for that data point to be visible.

### Evaluation of anti-E2 serological responses by ELISA

Four groups of mice (n=6 per group) were immunized with CHO sE2, geCHO.sE2.1, geCHO.sE2.7, and HEK sE2, which were formulated into nano-scale size supramolecular assemblies with a polyphosphazene adjuvant (PCPP-R) ^60–62^. Blood samples were collected prior to each vaccination on days 0 (pre-bleed), 14, 28, and 42 with a terminal bleed on day 56. Day 56 serum samples from the four groups of mice were individually tested for anti-E2 antibody titers in which the ELISA plates were coated with HEK sE2 (**Figure 5**). As shown, sera from mice immunized with any of the four antigens were able to induce an anti-E2-specific response of roughly equivalent magnitude. In addition, we performed an isotype-specific ELISA to analyze the magnitude of the humoral and cellular immune responses in the vaccinated mice. As shown in **Figure 5B** and **C**, the magnitude of the antibody (IgG1) and cell-mediated (IgG2a) responses are indistinguishable for the four groups. This is not surprising as the groups were immunized with equivalent amounts of antigen under the same regimen; the only difference resides in the glycan content of each antigen.

**Figure 5.**
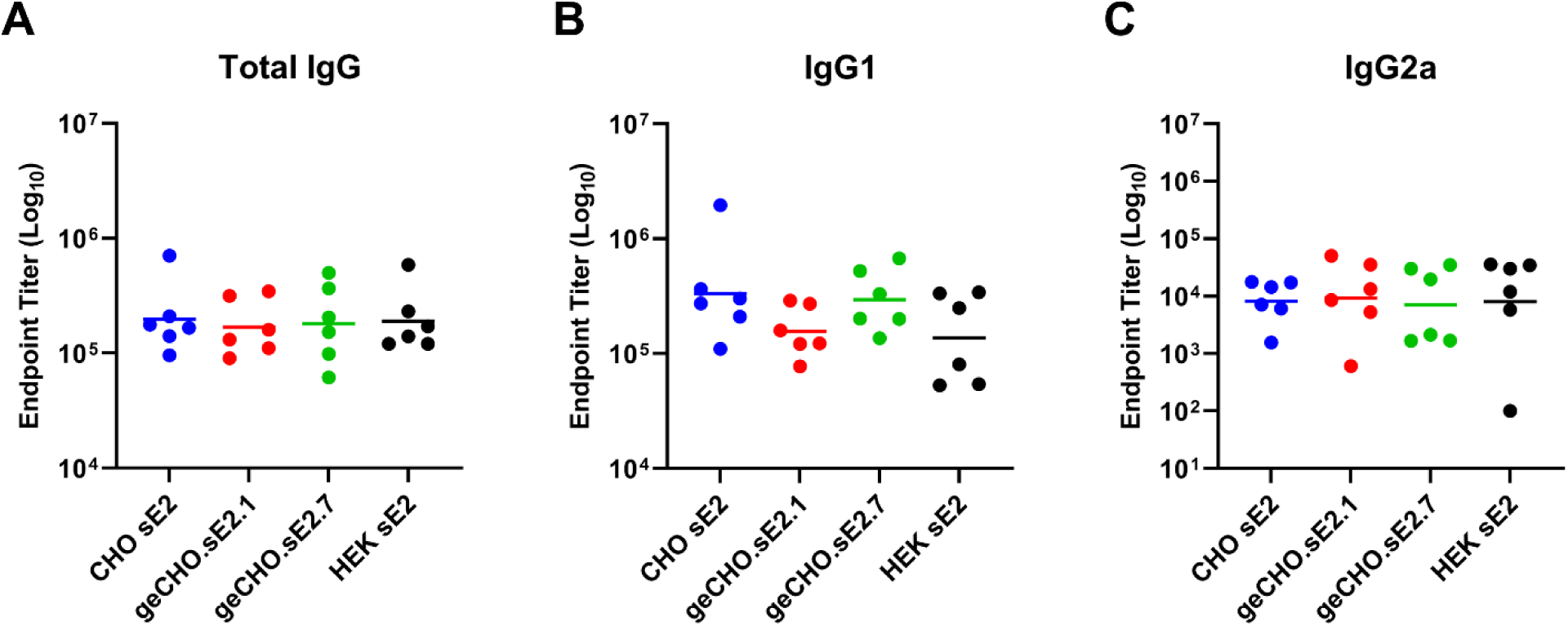
Assessment of antibodies induced in mice immunized with the four sE2 glycoforms at day 56 post-immunization by ELISA. (A) Total IgG endpoint titers. (B) IgG1 endpoint titers. (C) IgG2a endpoint titers. Endpoint titers were calculated by curve fitting in GraphPad Prism software with the endpoint OD defined as four times the highest absorbance value of day 0 sera.

### Evaluation of bnAb Responses by Competition Inhibition Analysis

The relative magnitude of domain-specific serological responses to conserved, continuous and discontinuous epitopes were analyzed by competition inhibition ELISA (**Figure 6**) using a panel of broadly neutralizing human monoclonal antibodies (HMAbs) derived from HCV-infected individuals ^63–68^. Purified IgGs from terminal bleed mouse antisera (day 56) from each group were used to compete with a non-nAb against domain A (CBH-4B) and HMAbs from the antigenic domains of the E2 ectodomain that give rise to bnAbs: domain B (AR3A/HEPC74), domain D (HC84.26.WH.5DL/HC84.1), and domain E (HCV1). There are no statistically significant differences between the groups for competition to any of the antibodies. For bnAbs AR3A (domain B) and HC84.26, there is robust competition for all groups, indicating an equivalent abundance of those types of antibodies in the immunized mouse sera for all groups. For HC84.1, the mean percent inhibition for each group are similar, with the HEK sE2 group being the lowest due to serum from one mouse exhibiting no competition for that bnAb and the geCHO.sE2.7 group exhibiting the most intra-group spread. However, there are some differences in antibody competition evident between the groups for CBH4B, HEPC74, and HCV1. In particular, the geCHO.sE2.7 group exhibits the highest level of competition for the non-nAb CBH4B and the lowest level of competition for the bnAb HEPC74. This might indicate a modest skewing of the immune response for mice vaccinated with the geCHO.sE2.7 antigen in an unfavorable direction. In contrast, the geCHO.sE2.1 group exhibited the highest level of competition for the bnAb HCV1 and comparably robust competition for the other bnAbs relative to the HEK and CHO sE2 groups. These differences suggest an influence of glycan content on the development of the immune response that warrants a more comprehensive exploration.

**Figure 6.**
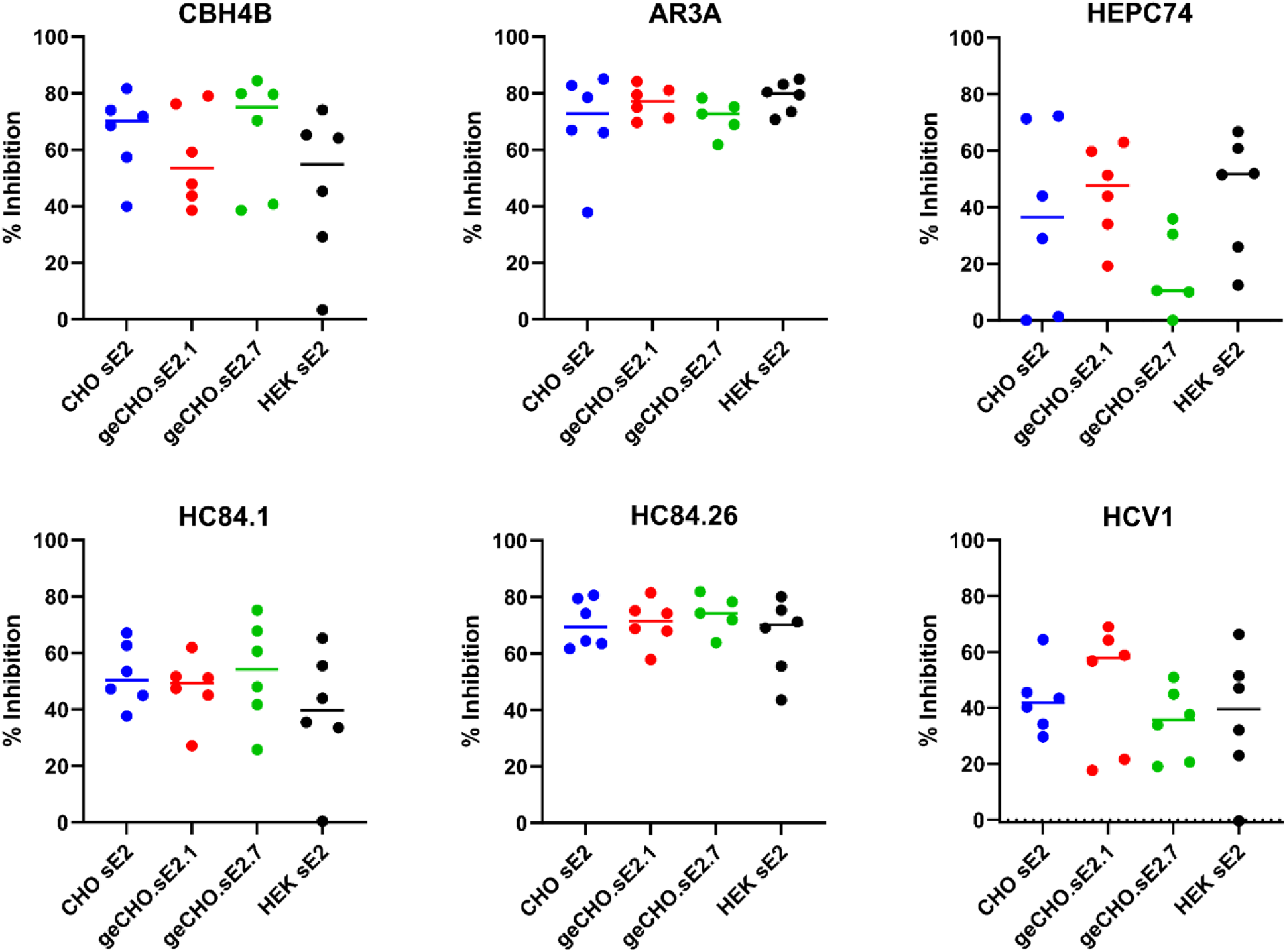
Competition ELISA of purified IgGs from individual mouse sera from day 56 after immunization. IgGs were diluted to a final concentration of 100 μg/mL and used to compete for binding to antibodies against antigenic domains A (CBH4B), B (AR3A and HEPC74), D (HC84.1 and HC84.26), and E (HCV1).

### Induction of bnAb responses

The ability of sera from mice immunized with CHO sE2, geCHO.sE2.1, geCHO.sE2.7, and HEK sE2 to inhibit HCV infection *in vitro* was tested against a panel of HCVpp covering the structural proteins of representative strains of the HCV genotypes responsible for the vast majority of infections in the Americas, Europe, and Asia (genotypes 1, 2, and 3). HCVpp packaged with the E1E2 glycoproteins of three antigenically distinct HCV genotypes (GTs), GT1a (H77C, AF011751), GT1b (UKNP1.18.1), GT2a (J6), GT2b (UKNP2.5.1), and GT3 (UKNP3.2.2) with two subtypes for both GT1 and GT2 were produced in HEK293T cells as described ^19^ and used for neutralization assays (**Figure 7**). Pre-immune and day 56 serum samples were used at two-fold serial dilutions starting at 1:64 (**Figure S7**) and inhibition values are expressed as the serum dilution level corresponding to 50% neutralization (ID_50_). Neutralization was most robust against the homologous genotype 1a strain (**Figure 7A**) as expected, with two notable differences observed among the groups. First, for the geCHO groups, the neutralization ID_50_ values exhibited much less spread among the mice within the group than for the CHO and HEK groups for GT1a neutralization. This suggests that increasing the homogeneity of glycan content reduces variability in bnAb response for this antigen. Second, serum from the geCHO.sE2.1 group is more effective at neutralizing the homologous H77 HCVpp than the other groups. The observed differences were not statistically significant, perhaps due to the large variability in ID_50_ for the CHO and HEK groups. However, the mean ID_50_ for the geCHO.sE2.1 group is 7-fold higher than for the CHO and HEK groups (**Table S2**) and 4-fold higher than the geCHO.sE2.7 group, indicating a clear increase in neutralization potency. Similarly, purified IgGs from the geCHO.sE2.1 group exhibited the most potent neutralization activity against an intergenotypic chimera cell culture-derived HCV (HCVcc, H77/JFH1, described in Materials and Methods) (**Figure 7B**). Cross-neutralization of heterologous strains was generally weak among all four groups, but an analysis of how many mice exhibited measurable heterologous neutralization activity for each of the groups shows that the geCHO.sE2.1 group had the most neutralization activity for all four heterologous HCVpp isolates tested (**Figure 7C**). Surprisingly, the geCHO.sE2.7 group performed the worst in neutralizing heterologous strains, with only one mouse exhibiting neutralizing activity for genotypes UKNP1.18.1 and J6 and lacking any mice with neutralizing activity for genotypes UKNP2.5.1 and UKNP3.2.2. The CHO and HEK groups had numbers of mice with measurable neutralizing activity intermediate between the geCHO.sE2.1 and geCHO.sE2.7 groups.

**Figure 7.**
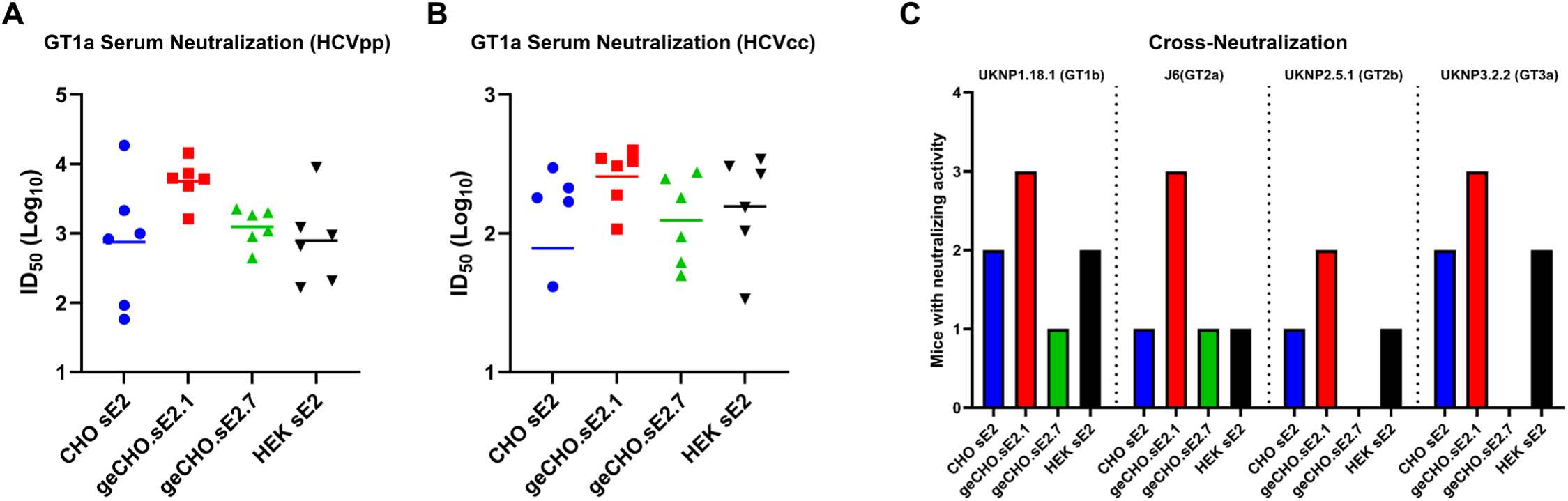
Improved neutralization by antibodies obtained from glycoengineered sE2. (A) Antibodies from the geCHO.1 line showed a 7-fold increased neutralization potency over antibodies from mice treated with the WT CHO and HEK cells for GT1a HCVpp. (B) Antibodies from the geCHO.1 line showed increased neutralization potency over antibodies from mice treated with the WT CHO and HEK cells for GT1a HCVcc. (C) Purified IgGs from the geCHO.1 line exhibited a higher frequency of cross-neutralization against HCVpp from genotypes 1b, 2a, 2b, and 3a.

### Analysis of the impact of glycan structure on serological responses

Based on the above data showing that distinct glycoforms of E2 vaccine candidates elicited different antibody responses, we further investigated which glycan structures were associated with optimal antibody-based neutralization. Taking into account the abundance of all of the glycoforms present in all of the antigens used in this study, we identified six glycans that significantly correlate with antibody IC_50_ values (**Figure 8**). Four glycans (masses 1375, 1620, 1794, 1824 Da) show a significant negative correlation (Spearman’s correlation: r=-0.46, p<0.05) with IC_50_s for mAb binding to H77C. We found that resulting antibodies with lower IC_50_ values and thus higher neutralization potential were obtained only for smaller glycans (i.e., biantennary, monoantennary) with terminal galactose or mannose. Moreover, the glycan with a mass of 1794 is the main glycan structure expressed in the geCHO.sE2.1, in which it showed a negative correlation (Spearman’s correlation: r<-0.15) with IC_50_s for all antibody binding. One glycan (2244) showed a significant negative correlation with IC_50_s for antibodies isolated from vaccinated animals and used against UKNP2.5.1. Meanwhile, the bi-antennary glycan with terminal GlcNAc (mass 1661) showed a positive correlation against all genotypes, especially significant for UKNP2.5.1 (Spearman’s correlation: r=0.53, p<0.01), suggesting that this glycan may lead to a higher IC_50_, corresponding to reduced serum binding and thus less potent neutralization, and should be avoided. Interestingly, this suggests that increased exposure to the underlying sE2 protein by decreased glycan size is not the only factor influencing sE2 antigenicity.

**Figure 8.**
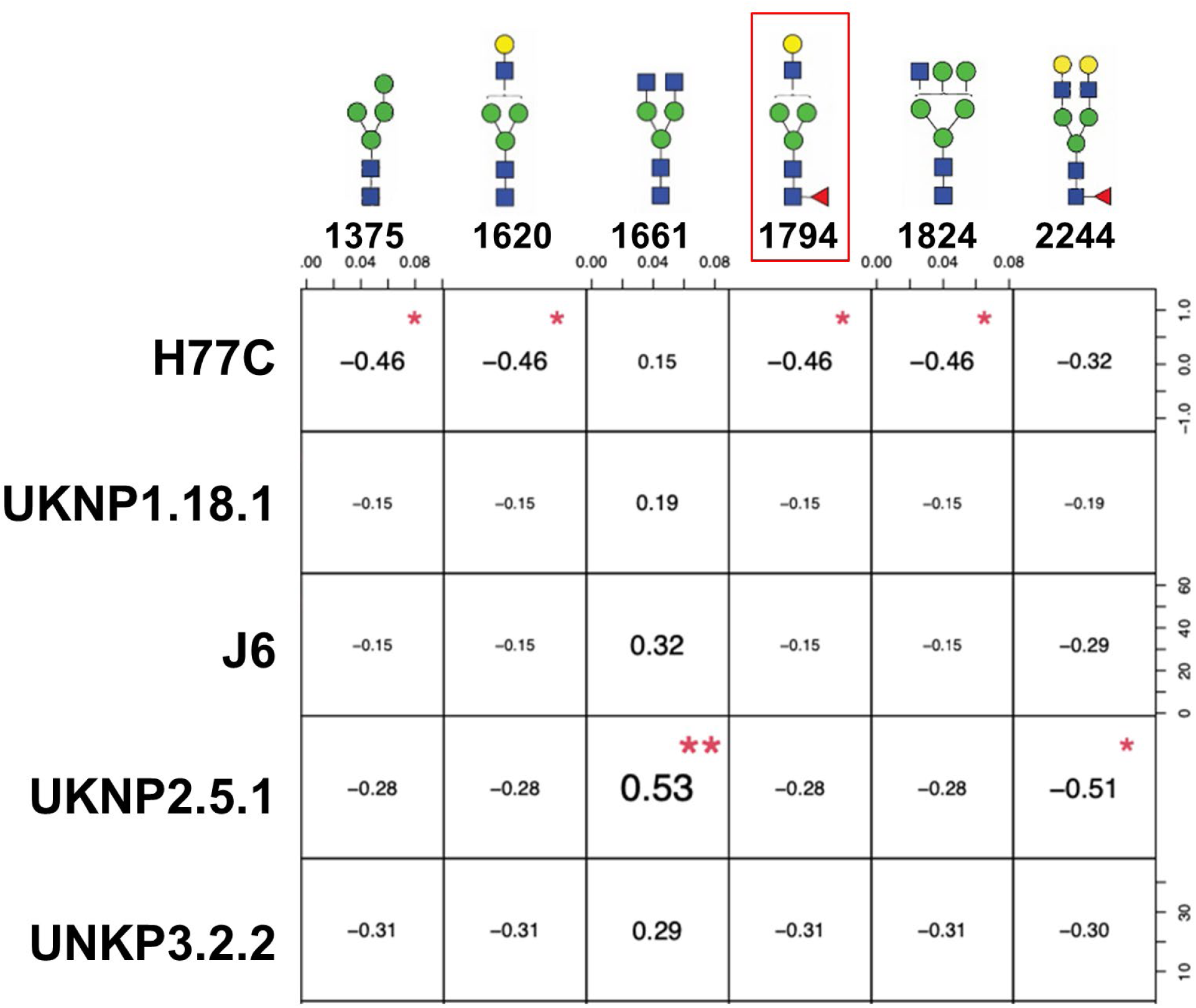
The correlation of glycan abundance on sE2 glycoforms and IC_50_ of antibodies generated in mice inoculated by the different glycoforms and used to treat different HCV genotypes. The correlation analysis conducted here is the Spearman rank correlation, and the numbers shown here are the Spearman’s rank correlation coefficient. Six glycans show significant correlations. The correlation coefficients are scaled relative to their P-value (i.e. the larger the number, the lower its associated P-value is).

We next computed the correlations between EPT and glycan structure (**Figure 9**). Six out of the seven identified glycans show a significant negative correlation (Spearman’s correlation: r<=-0.40, p<0.05) with EPT of IgG1, indicating that these glycans lead to the formation of more effective antibodies, seen in both the pooled IgGs and specifically IgG1. Interestingly, three of these six glycans were also in the set that correlated with lower IC_50_ values against the various HCV genotypes. Therefore, this finding confirms the need to focus on robust generation of an IgG1 response as part of HCV vaccine development.

**Figure 9.**
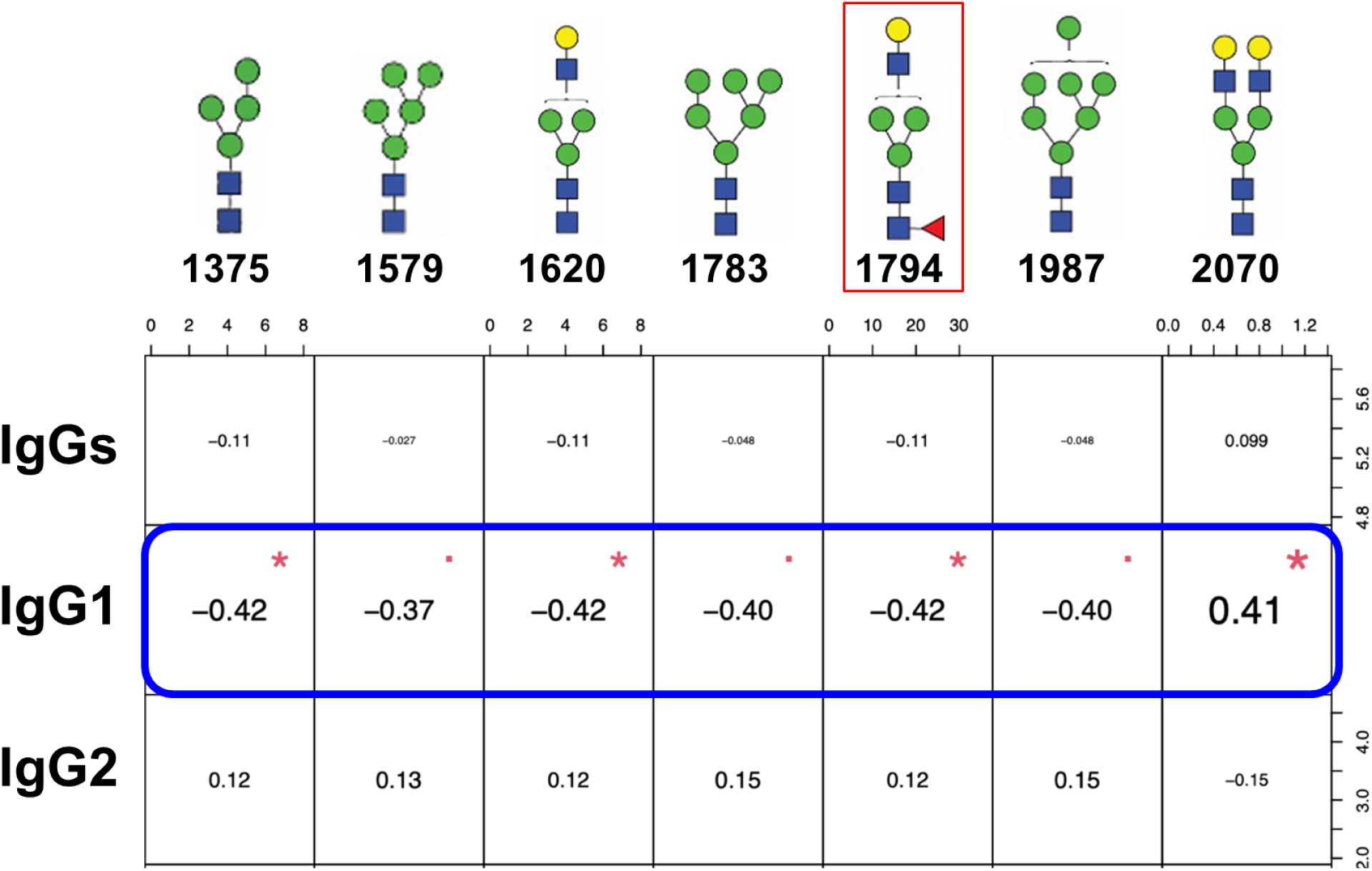
The correlation of glycan abundance on sE2 glycoforms and EPT of antibodies generated in mice inoculated by the different glycoforms and used to treat different HCV. The correlation analysis conducted here is the Spearman rank correlation, and the number shown here is the Spearman’s rank correlation coefficient. Seven glycans show significant correlations. The correlation coefficients are scaled relative to their P-value (i.e. the larger the number, the lower its associated P-value is).

## DISCUSSION

Most therapeutic proteins and viral vaccine antigens contain N-linked glycans that are important for stability and function. Since these glycans often shield epitopes from immune detection, both the extent to which a given vaccine antigen is glycosylated and the structure of the glycans imparted to that antigen will affect the immune response to the vaccine. Correspondingly, this will affect the immune memory response upon exposure to the natural pathogen to ward off infection. Ideally, the glycan content on a vaccine antigen would strike a balance between maximizing exposure to epitopes known to give rise to bnAbs while simultaneously avoiding the creation of neoepitopes due to exposure of regions that are not exposed to the immune system during a natural infection. At present, threading this needle is quite difficult due to limitations imposed by eukaryotic expression systems. Control of glycan content on recombinant proteins produced in off-the-shelf HEK, CHO, or insect cells is limited and thus the effect of different glycan structures on the immune response has not been explored in a systematic manner. Recently, our group developed geCHO cell lines that markedly increase the homogeneity of glycans imparted to recombinant proteins, thereby creating a platform with broad utility in the production of more precise biologic pharmaceuticals. To facilitate the study of glycan effects on immunogenicity, we leveraged these cell lines and investigated the effect of glycan content on the immune response to the HCV E2 ectodomain in mice. Our results represent a proof-in-principle demonstration that unique glycan structures on the HCV E2 glycoprotein impact immunogenicity and that analysis of biochemical and serological data can identify both beneficial and deleterious glycoforms.

Based on the biochemical data in this study, the chosen glycoforms did not appreciably impact solution behavior of the antigens. The geCHO sE2 antigens appeared to be more homogeneous by non-reducing western blot and AUC as expected due to the increased glycan homogeneity. This effect was more pronounced for geCHO.sE2.1, which has the smaller dominant glycan. One notable and surprising observation was that geCHO.sE2.7 exhibited poor binding to the CD81 receptor. The origins of this effect are unclear, but recent structural data for the E2-CD81 complex ^26^ might provide some clues. There are four glycans (N1, N2, N3, and N6) in the vicinity of the CD81 binding interface and, although none are visible in the electron density of the reported crystal structure, they could have an effect on binding. Alternately, this particular glycoform might attenuate the observed conformational changes in residues of antigenic domain E and the CD81 binding loop observed in the complex. The glycans N1, N2, and N3 are either in or adjacent to antigenic domain E and N6 resides on the CD81-binding loop (**Figure 10A, B**), which is consistent with either of the above potential effects. We also observed an effect on antigenicity and in particular for binding of antibodies AR3A and HEPC74 to antigenic domain B. This antigenic domain is bracketed by glycans N2, N4, N6, and N8 (**Figure 10C**) and in fact glycan-Fab contacts comprise approximately 23% of the E2-HEPC74 binding interface ^69^. While 23% is not likely determinative, the absence of these productive glycan-Fab interactions, or the presence of glycans that disfavor them, could impact HEPC74 binding and/or the development of HEPC74-like bnAbs during the course of the immune response upon vaccination. Since the presence of HEPC74-like bnAbs in patient sera correlates with spontaneous clearance ^28^, these results suggest that more attention needs to be paid to how glycan content affects both antigenicity and immunogenicity. That the observed effects on biochemical properties was modest (i.e. aside from receptor binding) is not surprising given the fact that the E2 ectodomain is generally well-behaved in solution and further that we selected for improved antigenicity and robust expression. It is likely that, had we performed a more comprehensive analysis, we would have found some glycans that negatively affected these parameters.

**Figure 10.**
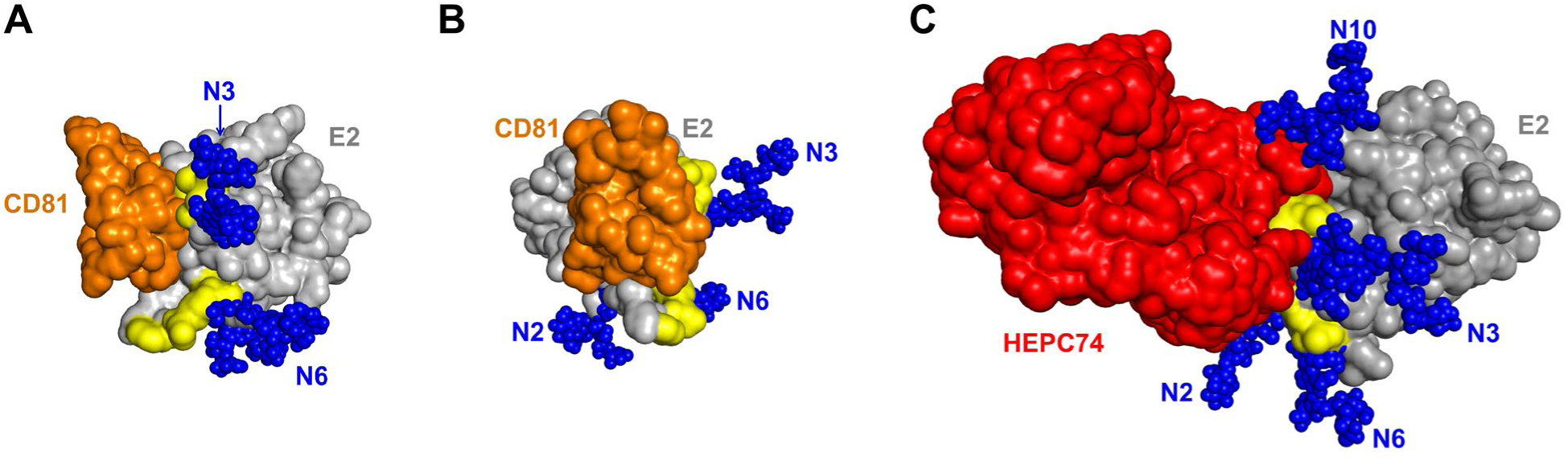
**A.** Three glycans (blue) in the E2 ectodomain are in close proximity the CD81 binding site. The CD81-LEL bound to E2 is shown as an orange surface. The three glycans at sites N2, N3, and N6, corresponding to attachment at residues N423, N430, and N532 (H77C numbering) respectively are shown as blue spheres. N2 is obscured in this orientation but visible in panel B. The E2 ectodomain is represented as a gray surface and antigenic domain B is represented as a yellow patch on the gray surface of E2. This model is adapted from the structure 7wmx ^26^. Because extended glycan chains were not resolved in the structure, a representative high mannose glycan was placed at each site to better show the extent of glycan shielding in the E2-CD81 complex. **B.** The same complex as in A, rotated 90 degrees relative to an axis running parallel to the screen. **C.** Four glycans (blue) in the E2 ectodomain are in close proximity to antigenic domain B. HEPC74 Fab bound to E2 is shown as a red solid surface. The four glycans at sites N2, N3, N6, and N10, corresponding to attachment at residues N423, N430, N532, and N623 (H77C numbering) respectively are shown as blue spheres. The E2 ectodomain is represented as a gray surface and antigenic domain B is represented as a yellow patch on the gray surface of E2. This model is adapted from the structure 8fsj ^69^. Because extended glycan chains were not resolved in the structure, a representative high mannose glycan was placed at each site to better show the extent of glycan shielding in antigenic domain B. The orientation matches that in panel A.

Examining the serological data shows that antibody titers elicited by the four antigens are approximately equivalent for total IgGs as well as the IgG1 and IgG2a isotypes. Similarly, the antibody competition assays reveal only modest differences, indicating that the distribution of antibodies to the different antigenic domains is largely the same. The response elicited by geCHO.sE2.7 contains proportionally more CBH4B-like antibodies and less HEPC74-like antibodies but these differences are not statistically significant. However, we do observe a pronounced difference in the neutralization potency of the immune response for the geCHO.sE2.1 group relative to the other three groups. The neutralization ID_50_ values are better for homologous pseudovirus neutralization and this group had more mice whose sera possessed neutralizing activity against the each of the heterologous pseudoviruses. Moreover, this neutralization profile was also observed for cell culture virus (HCVcc) derived from the hepatoma cell line, Huh7, that should more closely mimic the glycan profile of native virus. Thus, there is an improvement in the quality of the immune response for the geCHO.sE2.1 group that is not reflected in the antibody titers or the competition data when considered on their own. However, a deeper analysis that includes the glycan abundances for all antigens and the IC_50_ values for neutralization by purified IgG from mouse sera from the four groups revealed additional information about glycan effects on neutralization. In particular, there were six glycans that significantly correlated with better IC_50_ values, five of which were beneficial and one deleterious (**Figure 8**). Glycan profiling showed that geCHO.sE2.1 possessed all five of the beneficial glycans at >5% abundance and the deleterious glycan (mass 1661) at approximately 0.1% abundance. By contrast, geCHO.sE2.7, which elicited the weakest cross-neutralizing response, did not possess any of the five beneficial glycans in excess of 1% of the total population, but had the mass 1661 glycan at approximately 6% abundance. The WT CHO and HEK antigens had intermediate abundances, with WT CHO having more of the beneficial glycans than the HEK antigens (but fewer than geCHO.sE2.1) and also less of the mass 1661 glycan. However, these antigens also had many other glycans at significant abundance that did not correlate one way or the other with IC_50_, most likely due to the limited data set of this study that in a more comprehensive analysis might have shown effects not captured here. This study shows that a comprehensive analysis of biochemical properties, serological data, and glycan profiles can yield information about which glycans to retain and which ones to avoid for a given antigen. Such data can be used to choose a production host or growth conditions that favor beneficial glycans and suppress deleterious ones. The success of this small study indicates that a more comprehensive analysis can yield additional data regarding the potential effects of glycans on the efficacy of vaccine antigens. A more thorough, systematic analysis using all of the geCHO cell lines could reveal additional glycans that correlate with neutralization potency, along with epitope-specific effects of different glycans, perturbations to biochemical properties and changes to antigen structure. These data can then be used to guide vaccine design and production as part of an overall vaccine optimization program.

## METHODS

### Protein expression and purification

The clone for expressing the recombinant HCV sE2 has been described previously ^15, 18, 55^. Expression of CHO sE2 and geCHO.sE2.1 and geCHO.sE2.7 was performed as follows. DNA encoding sE2 was chemically transfected using Fugene (Promega, Madison, WI) in CHO cultures after which the transfected cells were allowed to express and secrete sE2 into the medium for 4-5 days. The supernatants were then harvested via centrifugation and stored at −80 °C prior to purification. Expression of recombinant HEK sE2 was performed via transient expression in human Expi293 cells using the Expi293 Expression System by following the manufacturer’s protocols (Thermo Fisher Scientific, Walthan, MA). Briefly, Expi293 cells were cultured in Expi293 Expression Medium in the shaker incubator at 37 °C, with 120 rpm and 8% CO_50_. When the cells reached a density of 2.0 × 10^6^ cells/mL, Expi293 cells were transfected using proper amounts of plasmid DNA. Culture supernatants of HEK sE2 were harvested at 72 hours after transfection, clarified by centrifugation at 10,000 rpm for 10 min, and filtered by a 0.22 μm filters. All four antigens were purified using the same protocol, which entailed subjecting the clarified supernatant to sequential HisTrap Ni^2+^-NTA and Superdex 200 size exclusion chromatography (SEC) as described in our previous papers ^15, 70^.

### SDS-PAGE and western blot

Purified sE2 antigens were separated by a precast, 4–20% Mini-PROTEAN TGX stain-free gels on a Mini-PROTEAN Tetra cell electrophoresis instrument (Bio-Rad Laboratories, Hercules, CA). In reducing conditions, each sample was incubated with loading dye (4x Laemmli buffer + 10% β-mercaptoethanol) (Bio-Rad Laboratories, Hercules, CA) and heated to 95 °C. In non-reducing conditions, each sample was incubated with Laemmli buffer and heated to 37 °C. For western blot detection, the purified protein samples on SDS-PAGE were transferred onto Trans-Blot Turbo Mini nitrocellulose membranes (Bio-Rad Laboratories, Hercules, CA). The membranes were then probed using the anti-HCV E2 mAb HCV1^67^ (purified in-house) at 5 μg/mL followed by detection using a secondary goat anti-human IgG-HRP conjugate (Invitrogen, Waltham, MA) at a 0.16 μg/mL dilution and the Western ECL substrate (Bio-Rad Laboratories, Hercules, CA). All gels were imaged using the ChemiDoc system (Bio-Rad Laboratories, Hercules, CA).

### N-linked glycomics analysis by mass spectrometry

Protein samples were reduced with dithiothreitol (DTT) at 50 °C for 30 minutes and subsequently desalted using a 3 kDa molecular weight cutoff (MWCO) filter with 50 mM ammonium bicarbonate buffer. Protein concentration was determined using a BCA assay (Thermo Fisher Scientific, Waltham, MA). For N-glycan release, equal amounts of protein were treated with PNGase F (New England Biolabs, Ipswich, MA) at 37 °C for 48 hours. Released N-glycans were collected using a 10 kDa MWCO spin filter (MilliporeSigma, Burlington, MA), followed by C18 cartridge purification, and lyophilized. N-linked glycans were permethylated using methyl iodide in the presence of NaOH-DMSO, as previously described ^71^. Permethylated glycans were analyzed using mass spectrometry techniques. For MALDI-TOF/TOF MS, glycans were mixed with 2,5-dihydroxybenzoic acid (DHB) matrix and analyzed on a RapifleX® MALDI-TOF/TOF MS (Bruker, Billerica, MA) instrument in reflector mode. For ESI-MS, glycans were dissolved in methanol:water (1:1) and analyzed using direct infusion mode on an Orbitrap Fusion Tribrid (Thermo Fisher Scientific, Waltham, MA) in positive ion mode. Data were collected from 200 to 2000 m/z and deconvoluted using FreeStyle 1.8 SP1 software (Thermo Fisher Scientific, Waltham, MA). Peak mass lists from both MALDI-MS and ESI-MS were analyzed using GlycoWorkbench software. Glycan relative abundance was calculated for each sample based on peak area.

### Analytical Ultracentrifugation

Sedimentation velocity (SV) experiments were performed at 20 °C using a ProteomeLab Beckman XL-A with absorbance optical system and a 4-hole An60-Ti rotor (Beckman Coulter, Indianapolis, IN). For each sE2 glycovariant, the sample and reference sectors of the dual-sector charcoal-filled epon centerpieces were loaded with 380 μL protein in PBS, pH 7.4, and 400 μL buffer. The cells were centrifuged at 40 krpm and the absorbance data were collected at 280 nm in a continuous mode with a step size of 0.003 cm and a single reading per step to obtain linear signals of <1.25 absorbance units. Sedimentation coefficients were calculated from SV profiles using the program SEDFIT ^72^. The continuous *c*(*s*) distributions were calculated assuming a direct sedimentation boundary model with maximum entry regularization at a confidence level of 1 standard deviation. The density and viscosity of buffers at 20 °C and 4 °C were calculated using SEDNTERP ^73^. The *c*(*s*) distribution profiles were prepared with the program GUSSI (C.A. Brautigam, Univ. of Texas Southwestern Medical Center).

### Enzyme-linked immunosorbent assay (ELISA) screening of sE2 glycovariants for mAb and receptor binding

HCV HMAb binding to the thirteen glycovariants of sE2 were evaluated and quantitated by ELISA. 96-well half-area microplates (Grenier Bio-One, Thermo Fisher Scientific, Waltham, MA) were coated with 5 μg/mL Galanthus Nivalis Lectin (Vector Laboratories, Newark, CA) overnight, and purified sE2 was then added to the plates at 2 μg/ml. After the plates were washed with PBS and 0.05% Tween 20, and blocked by Pierce™ Protein-Free (PBS) Blocking Buffer (Thermo Fisher Scientific, Waltham, MA), the mAbs were tested in duplicate at 3-fold serial dilution starting at 20 nM. The total volume for these experiments was 35 μL per well owing to the limited amounts for some samples. For receptor binding and additional antibody-binding measurements 96-well microplates (MaxiSorp, Thermo Fisher Scientific, Waltham, MA) with a final volume of 100 μL per well were used. For receptor binding experiments, a CD81 large extracellular loop (LEL)-Fc fusion (kind gift of Dr. Mansun Law, Scripps Research) was used at a starting concentration of 2.8 μM and serially diluted two-fold. These experiments were conducted in duplicate. The binding was detected by HRP-conjugated anti-human IgG secondary antibody (Invitrogen, Waltham, MA) at a concentration of 0.16 mg/mL with TMB substrate (Bio-Rad Laboratories, Hercules, CA). The absorbance was read at 450 nm using a SpectraMax MS microplate reader (Molecular Devices, San Jose, CA). For detection of antibody binding, the same procedure was employed with the exception that the starting antibody concentration was 66 nM. The data were analyzed by nonlinear regression to measure antibody dissociation constants (K_d_) using GraphPad Prism software.

### Animal immunization

CD1 mice were purchased from Charles River Laboratories. Prior to immunization, sE2 glycovariants were formulated with polyphosphazene adjuvant as described in previous studies ^60,62^. In brief, 50 μg poly[di(carboxylatophenoxy)phosphazene (PCPP) was formulated with 25 μg resiquimod, R848 in PBS (pH 7.4) to form the PCPP-R adjuvant. The resulting supramolecular complex (PCPP-R) was formulated with sE2 antigen (50 μg for prime or 10 μg for boost immunization). Dynamic light scattering (DLS) was used to confirm the absence of aggregation in adjuvanted formulations. Groups of six female CD-1 mice, age 7 to 9 weeks, were immunized via the intraperitoneal (IP) route, first with a prime as described above on day 0, then with boosts as described above on day 14, day 28 and day 42. Blood samples were collected prior to each vaccination on days 0 (pre-bleed), 14, 28, 42 and a terminal bleeding on day 56. The blood samples were processed for serum by centrifugation and stored at −80 °C until analysis was performed.

### ELISAs for serum antibody detection

ELISA was performed to measure HCV E2-specific antibody responses in sera from immunized mice. 96-well plates (MaxiSorp, Thermo Fisher Scientific, Waltham, MA) were coated overnight with 5 µg/mL Galanthus Nivalis Lectin (Vector Laboratories, Newark, CA) at 4 °C. The next day, plates were washed with PBS containing 0.05% Tween 20 and coated with 200 ng/well of the sE2 glycovariants at 4 °C. After overnight incubation, plates were washed with PBS containing 0.05% Tween 20 and blocked with Pierce™ Protein-Free Blocking Buffer (Thermo Fisher Scientific, Waltham, MA) for 1 hour, and serially diluted mice sera samples were then added to the plates and incubated for another hour. The binding of HCV E2-specific antibodies was detected by an HRP-conjugated anti-mouse IgG secondary antibody (Abcam, Waltham, MA) at a concentration of 0.4 μg/mL with TMB substrates (Bio-Rad Laboratories, Hercules, CA). Absorbance values at 450 nm (SpectraMax M3 microplate reader) were used to determine endpoint titers, which were calculated by curve fitting in GraphPad Prism software and defined as four times the highest absorbance value of pre-immune sera. Statistical analysis to determine significance was performed using a Kruskal-Wallis one-way ANOVA.

### Competition ELISA

The ability of antibodies in immunized mouse sera to compete with conformation-dependent or linear HCV E2-specific HMAbs was assessed by ELISA. The antibodies used for these experiments include AR3A ^56^ and HEPC74 ^74^ (domain B), HC84.26 ^57^ and HC84.1 ^75^ (domain D), HCV1 ^67^ (domain E), and CBH-4B ^76^ (domain A). sE2 was captured via incubation on GNA-coated microtiter plates at 4 °C overnight. After blocking with Pierce™ Protein-Free Blocking Buffer (Thermo Fisher Scientific, Waltham, MA) for 1 hour followed by three washes using the same buffer, diluted IgGs purified from terminal bleed mouse antisera were added to each well and incubated for 1 hour at room temperature. After plates were washed with PBS containing 0.05% Tween 20, HCV E2-specific HMAbs were added at a concentration demonstrated previously to result in 70% of maximal binding and incubated for an additional hour. The HMAbs used for the competition ELISA were biotinylated using an EZ-Link NHS-PEO solid-phase biotinylation kit (Thermo Fisher Scientific, Waltham, MA). Bound biotinylated HMAb was detected using HRP-conjugated streptavidin (Abcam, Waltham, MA) at a concentration of 0.05 μg/mL. Absorbance was read at 450 nm using a SpectraMax M3 microplate reader. Percent inhibition values were calculated as the percentage of mAb binding relative to the mAb bound in the absence of serum.

### HCVpp neutralization assay

Total serum IgGs were purified using protein G HP SpinTrap columns (Cytiva, Marlborough, MA) according to the manufacturer’s protocol. Briefly, 600 μL of mouse serum was loaded on the column and incubated for 4 min with gentle mixing. After washing 2 times with binding buffer (20 mM sodium phosphate, pH 7.0), IgGs were eluted with elution buffer (0.1 M glycine-HCl, pH 2.7) into tubes containing neutralizing buffer (1 M Tris-HCl, pH 9.0). For HCVpp neutralization, purified IgGs were serially diluted from a starting concentration of 100 μg/mL or 500 μg/mL.

The human hepatoma cell line, Huh7, was maintained in the DMEM medium supplemented with 10 % FBS and 1% non-essential amino acids (NEAA) (Thermo Fisher Scientific, Waltham, MA), and used as the target cell line for neutralization assays ^15, 19^. To test sera and antibodies for neutralization, Huh7 cells were pre-seeded into 96-wells plates at a density of 1 × 10^4^ per well. In next day, the pseudoparticles were incubated with defined concentrations of mAbs and/or the heat-inactivated serum at indicated dilutions for 1 hour at 37 °C, and then added to each well. After the plates were incubated in a CO_50_ incubator at 37 °C for 5 to 6 hours, the mixtures were replaced with fresh medium and then continued to incubate for 72 hours. After incubation, 100 μl of Bright-Glo reagent (Promega, Madison, WI) was added to each well for 2 minutes at room temperature and the luciferase activity was measured using a FLUOstar Omega plate reader (BMG Labtech, Cary, NC) with the MARS software. The 50% inhibitory concentration (IC_50_) was calculated as the mAbs concentration that caused a 50% reduction in relative light units (RLU) compared with pseudoparticles in the control wells. Titers of nAbs in animal sera were reported as 50% inhibitory dilution (ID_50_) values. All values were calculated using a dose-response curve fit with nonlinear regression in GraphPad Prism. All experiments involving the use of pseudoparticles were performed under biosafety level 2 conditions.

### HCVcc generation and titration

HCV viral RNA was produced via *in vitro* transcription of an XbaI-linearized BiGluc-H77C(1a)/JFH (T2700C/A4080T ^77, 78^) (kindly provided by Jens Bukh, University of Kopenhagen) plasmid using the HiScribe® T7 High Yield RNA Synthesis Kit (New England Biolabs, Ipswich, MA, E2040S) as outlined in the user manual. Viral RNA was purified using the MEGAclear™ Transcription Clean-Up Kit (Thermo Fisher Scientific, Waltham, MA AM1908) following manufacturer’s instructions, and quality control was performed by gel electrophoresis to ensure no significant RNA degradation. Viral RNA stocks were stored as 5 µg aliquots at −80 °C. RNA was electroporated into Huh7.5-1 cells (kindly provided by Frank Chisari, The Scripps Research Institute). The pellet was resuspended in the appropriate volume of cold DPBS to achieve a concentration of 1.5E7 cells/mL. 6E6 cells were then electroporated in a 2 mm path length electroporation cuvette (BTX Harvard Apparatus; Holliston, MA) with 5 µg of viral RNA using an ECM 830 Square Wave Electroporation System (BTX) at the following settings: five pulses, 99 µs per pulse, 1.1 s pulse intervals, 860V. Following a ten-minute incubation at room temperature, the electroporated cells were seeded into p150s and maintained in 5% FBS DMEM. Media was changed one day post-electroporation, and supernatants were collected twice daily from days four through six and stored at 4 °C. The pooled supernatants were passed through a 0.22 µm vacuum filter and subsequently concentrated to ∼40 mL in 100kDa MWCO Amicon® Ultra-15 Centrifugal Filter Units (MilliporeSigma, Burlington, MA, UFC9100).

The TCID_50_/mL of concentrated virus was determined after one freeze-thaw by limiting dilution assay. Huh7.5 cells (kindly provided by Charles Rice, The Rockefeller University) were seeded in a 96-well plate at a concentration of 6400 cells/well. 50 μL of ten-fold serial dilutions (from neat to 1:1E5) of virus were added to each column of wells, with 8 wells receiving each dilution. After removal of the inoculum 6-8 hours post-infection, cells were washed with unsupplemented DMEM and cultured in 200 μL DMEM containing 10% FBS and 1% penicillin/streptomycin. On day 3 post-infection, cells were fixed and permeabilized in ice-cold 100% methanol for 30 min at −20°C. Cells were blocked in 1X PBS containing 0.1% Tween-20, 1% BSA, and 0.2% skim milk for 30min at room temperature (RT). Cells were then treated with PBS containing 3% H_2_O_2_for 5 min at RT. Cells were then stained with a mouse anti-HCV NS5A monoclonal antibody (clone 9E10, 220 ng/mL, 50µL/well) for 1 hour at RT, followed by an HRP-conjugated goat anti-mouse polyclonal antibody (Invitrogen G-21040, 5 µg/mL, 50 µL/well). HRP signal was detected using DAB Peroxidase (HRP) Substrate Kit (Vector Laboratories, Newark, CA, SK4100). TCID_50_/mL was calculated using the Reed & Munch method ^79^.

### HCVcc neutralization assay

Huh7.5 cells (kindly provided by Charles Rice, The Rockefeller University) were seeded in 96-well plates at a concentration of 6400/well. Ten-fold serial dilutions were performed for pooled serum samples starting at 200 µg/mL, which were mixed 1:1 with HCVcc (final MOI = 0.1) and incubated for 1 hour at 4 °C. After incubation, serum and virus mixture was added to cells and cultured for 4 hours at 37 °C. After removal of the inoculum, cells were washed with twice with unsupplemented DMEM (Gibco, Waltham, MA) and then cultured in 100 μL DMEM containing 3% fetal bovine serum (FBS, Atlanta biologicals), nonessential amino acids (NEAA, 0.1 mM, Thermo Fisher Scientific, Waltham, MA), HEPES (20mM, Thermo Fisher Scientific, Waltham, MA), polybrene (4 μg/mL, Sigma-Aldrich Chemie GmbH, Burlington, MA) and penicillin streptomycin at 37 °C. After 72 hours, supernatants were collected and luciferase assay was performed following the manufacturer’s protocol (GeneCopoeia Inc.). Percent neutralization was calculated as relative luminescence units (RLU) from supernatant cultured without HCVcc (100% neutralization). The serum concentration of 50% neutralization (IC_50_) was calculated from the sigmoid curve (GraphPad Prism 10).

### Pair-wise correlation analysis between the glycans/epitopes and mAb binding strength

To determine pairwise correlation relationships between glycans (or epitopes) and mAb binding strength, we used measures of the Spearman’s rank correlation coefficient ^80, 81^, which describe the directionality and strength of the relationship between two studied variables. Specifically, we calculated pairwise Spearman’s rank correlation coefficients for the relative abundance of glycan structures or glycan epitopes with nAb binding.

## DATA AVAILABILITY

The data supporting the results of this study are included in the article/supplementary material. Further inquiries can be directed to the corresponding authors.

## ACKNOWLEDGEMENTS

We thank Yunus Abdul and Noble Life Sciences for performing the animal studies. Glycomics analysis was performed at the Complex Carbohydrate Research Center and was supported in part by the National Institutes of Health (NIH)-funded R24 grant (R24GM137782) to Parastoo Azadi. This work was also supported by NIAID (R01 AI132212, TRF, ET, AA; R01 AI168048, TRF, EAT, AA, AP; R01AI107301, AP), NIGMS (R35 GM119850, NEL; T32GM007388, MPS), and the Novo Nordisk Foundation (NNF20SA0066621, BGV, NEL).

## AUTHOR CONTRIBUTIONS

T.R.F., E.A.T., and N.E.L. developed the concept of the project. T.R.F., E.A.T., N.E.L., K.L.C., P.A., B.G.V., R.A.M., A.K.A., and A.P. designed the experiments. L.K., K.H.L., A.W.T.C., M.P.S., S.S., K.E., K.L.C., S.S., B.K., N.B.M., S. A-H., and A.M. conducted experiments. L.K., K.H.L., A.W.T.C., M.P.S., S.S., K.E., K.L.C., S.S., B.K., N.B.M., S. A-H., A.M., N.E.L., and E.A.T. performed data analysis. L.K., T.R.F., E.A.T., and N.E.L. wrote the manuscript. K.L.C., M.P.S., and A.P. provided substantive revisions. L.K., T.R.F., E.A.T., and N.E.L. revised and edited the manuscript. T.R.F coordinated the animal study. All authors approve of the submitted manuscript.

