## Supplemental Data for "Glycoengineering of the hepatitis C virus E2 glycoprotein leads to improved biochemical properties and enhanced immunogenicity"

*Corresponding author

**This PDF file includes:**

Supplementary Figures 1 to 7

Supplementary Tables 1 and 2

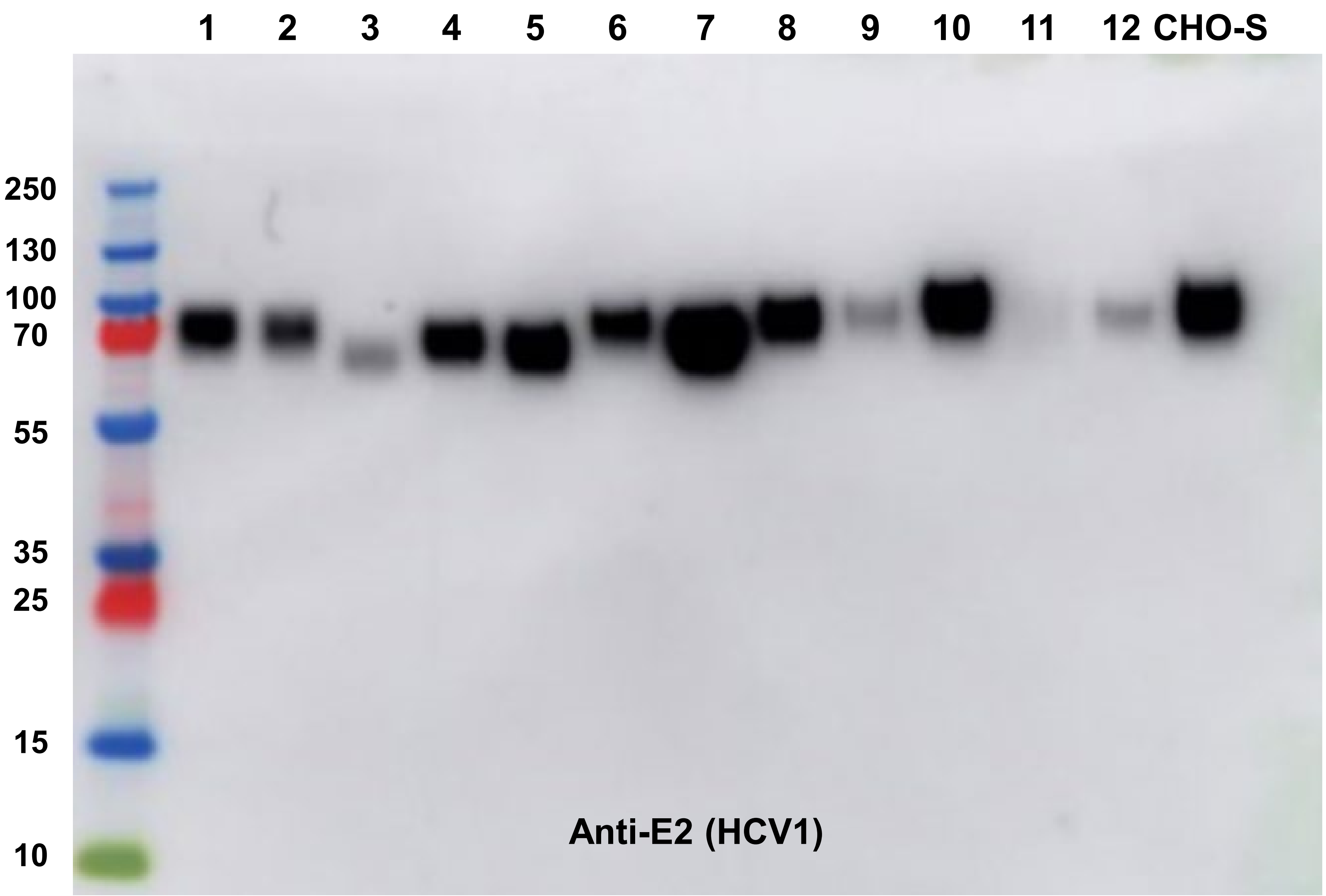


**Figure S1A.** Western blot analysis of sE2 expression in 12 geCHO cell lines and CHO-S. The anti-E2 antibody HCV1 antibody was used at a concentration of 5 μg/mL for the Western blot. Molecular weights, in kilodaltons, of the Western blot markers are indicated on the left.

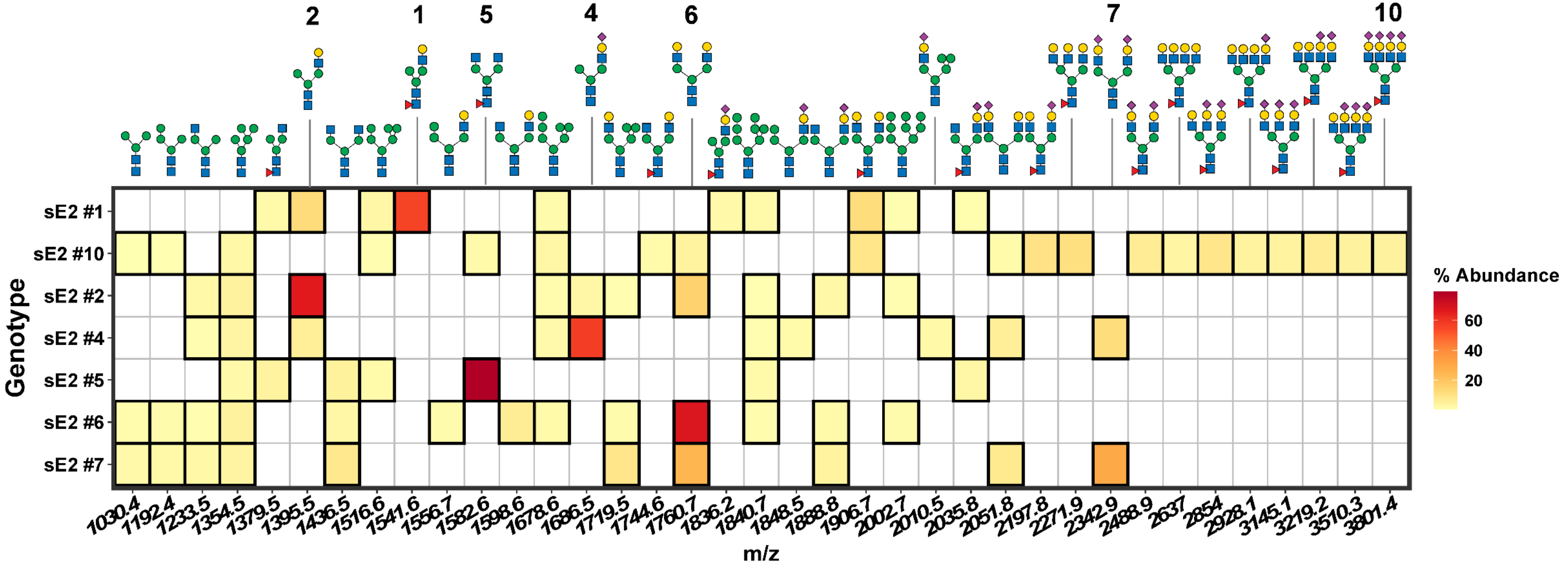


**Figure S1B.** The glycoprofiles of cells expressing in sE2. Heatmap representation of the relative abundance of N-linked glycans on recombinant sE2 in seven of the twelve glycoforms produced. The other five did not give glycan profiles of sufficient quality to include them in the heatmap due to insufficient amounts of protein produced. The most abundant glycan for each of the cell lines labeled with the cell line number above the corresponding glycan.

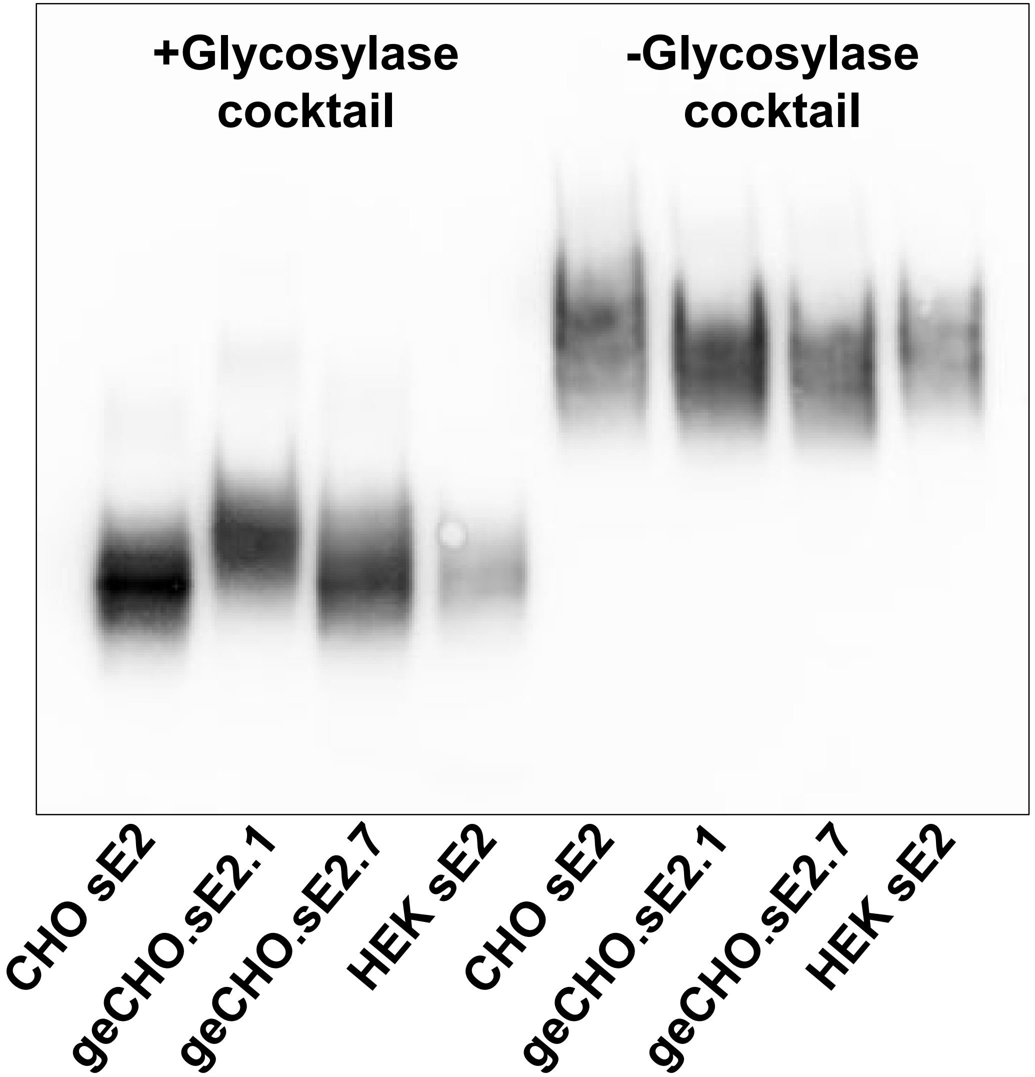


| Antigen | $K_d$ (WT HEK)/$K_d$ | | | | |
|---|---|---|---|---|---|
| | AR3A | HEPC74 | HC84.1 | HC84.26 | HCV1 |
| CHO sE2 | 0.9 | 1.0 | 0.9 | 0.7 | 1.4 |
| geCHO.sE2.1 | 0.8 | 0.8 | 0.4 | 0.4 | 1.0 |
| geCHO.sE2.7 | 0.6 | 0.7 | 0.5 | 0.4 | 0.8 |
| HEK sE2 | 1.0 | 1.0 | 1.0 | 1.0 | 1.0 |

**Figure S2.** Deglycosylation significantly attenuates glycoform-dependent differences in antibody binding. After incubation with a cocktail of glycosylases, the sE2 antigens shift to a more uniform migration. The antigen geCHO.sE2.1 appears to retain some glycan that the other antigens do not, as evidenced by slightly slower migration. The antigens after glycan removal exhibit smaller differences in antibody binding that the fully-glycosylated antigens in Table 1.

### CHO-S

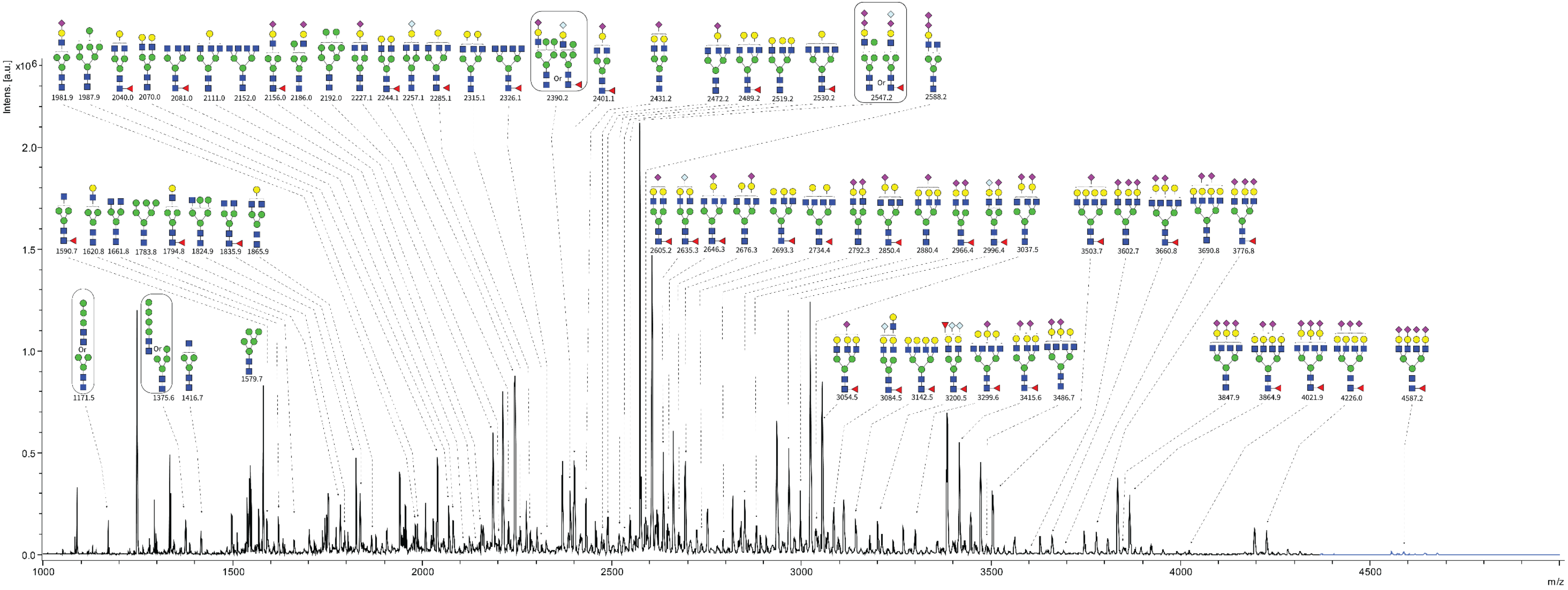


**Figure S3.** Glycan profile of sE2 produced in CHO-S cells.

### geCHO.1

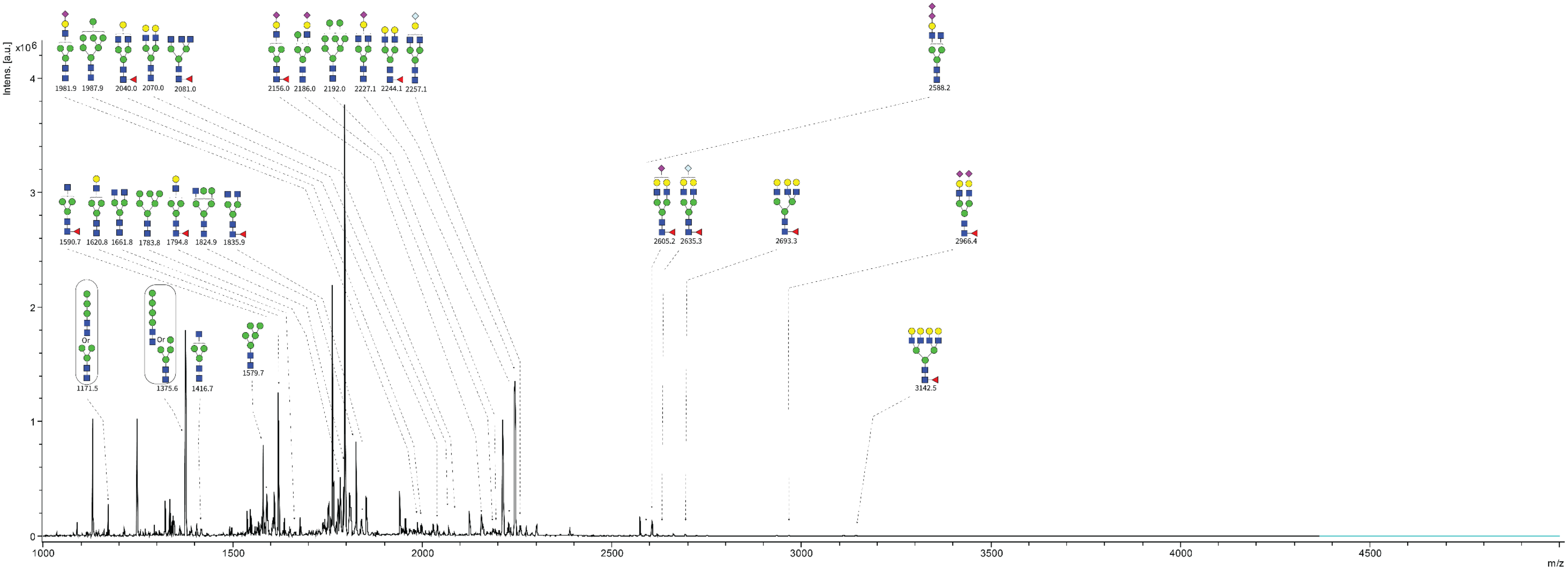


**Figure S4.** Glycan profile of sE2 produced in geCHO.1 cells.

### geCHO.7

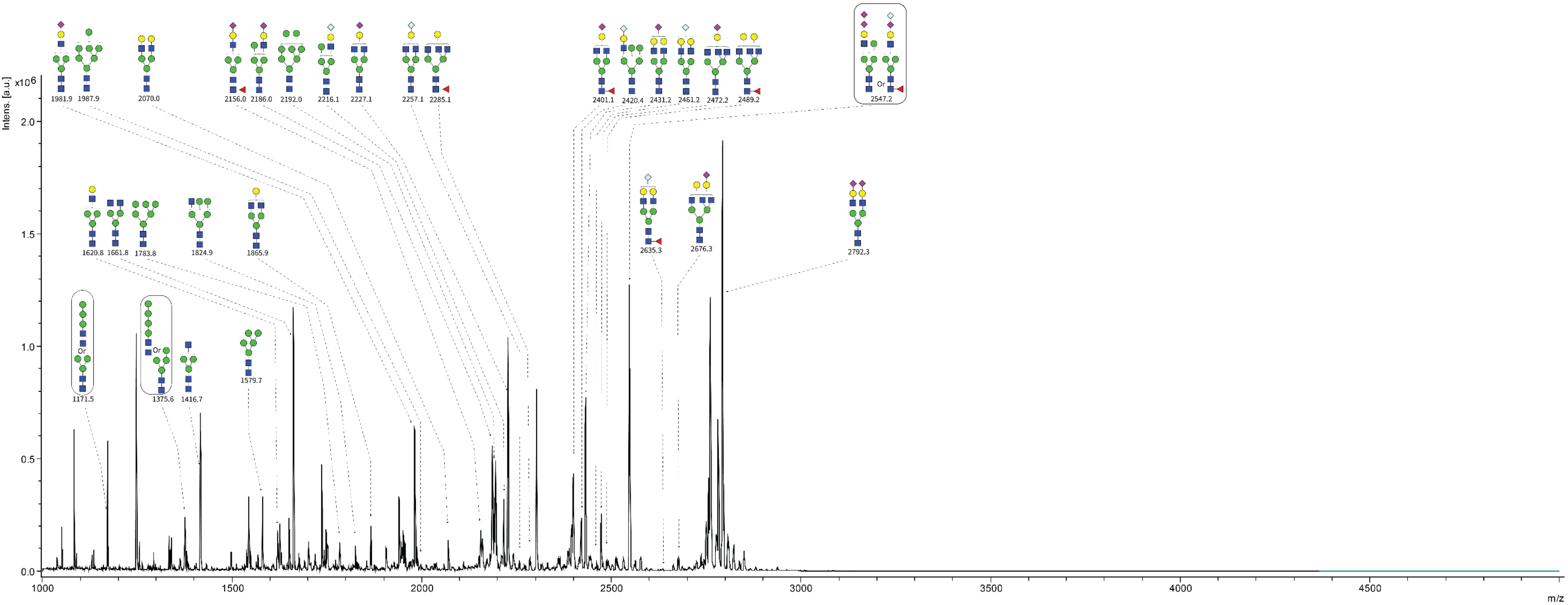


**Figure S5.** Glycan profile of sE2 produced in geCHO.7 cells.

### HEK 293

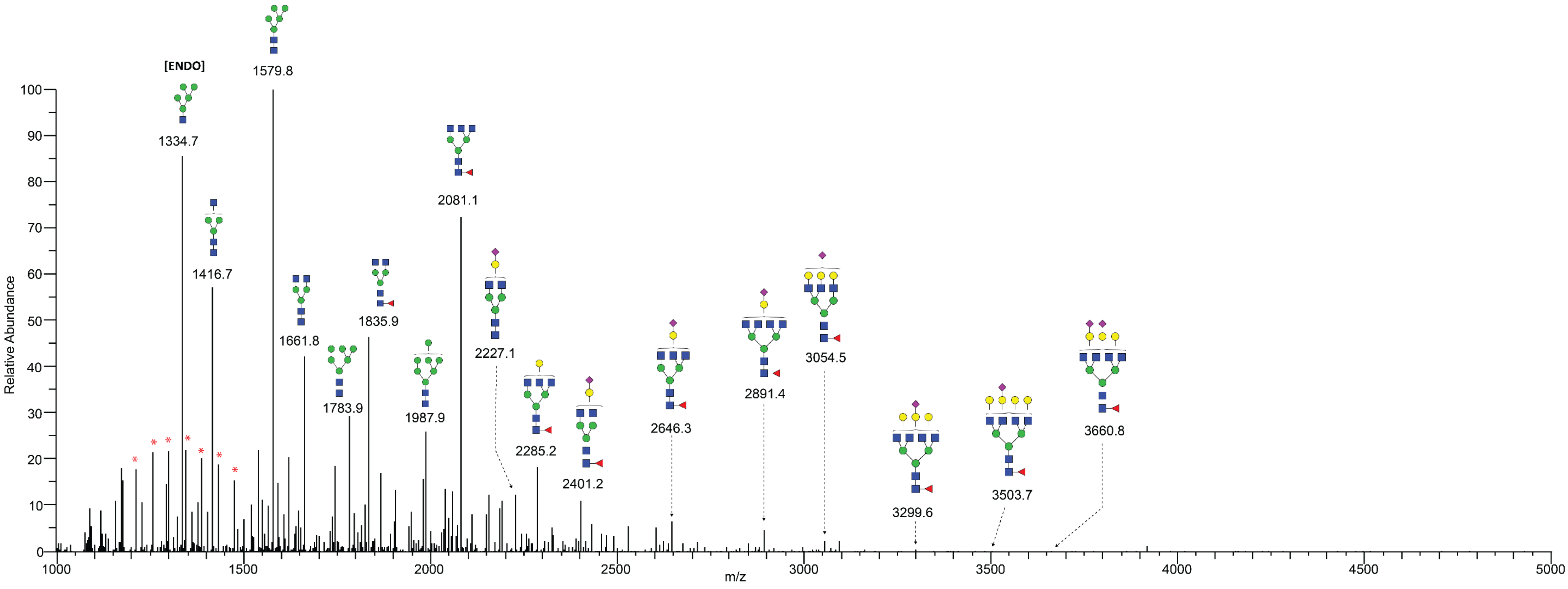


**Figure S6.** Glycan profile of sE2 produced in HEK 293 cells.

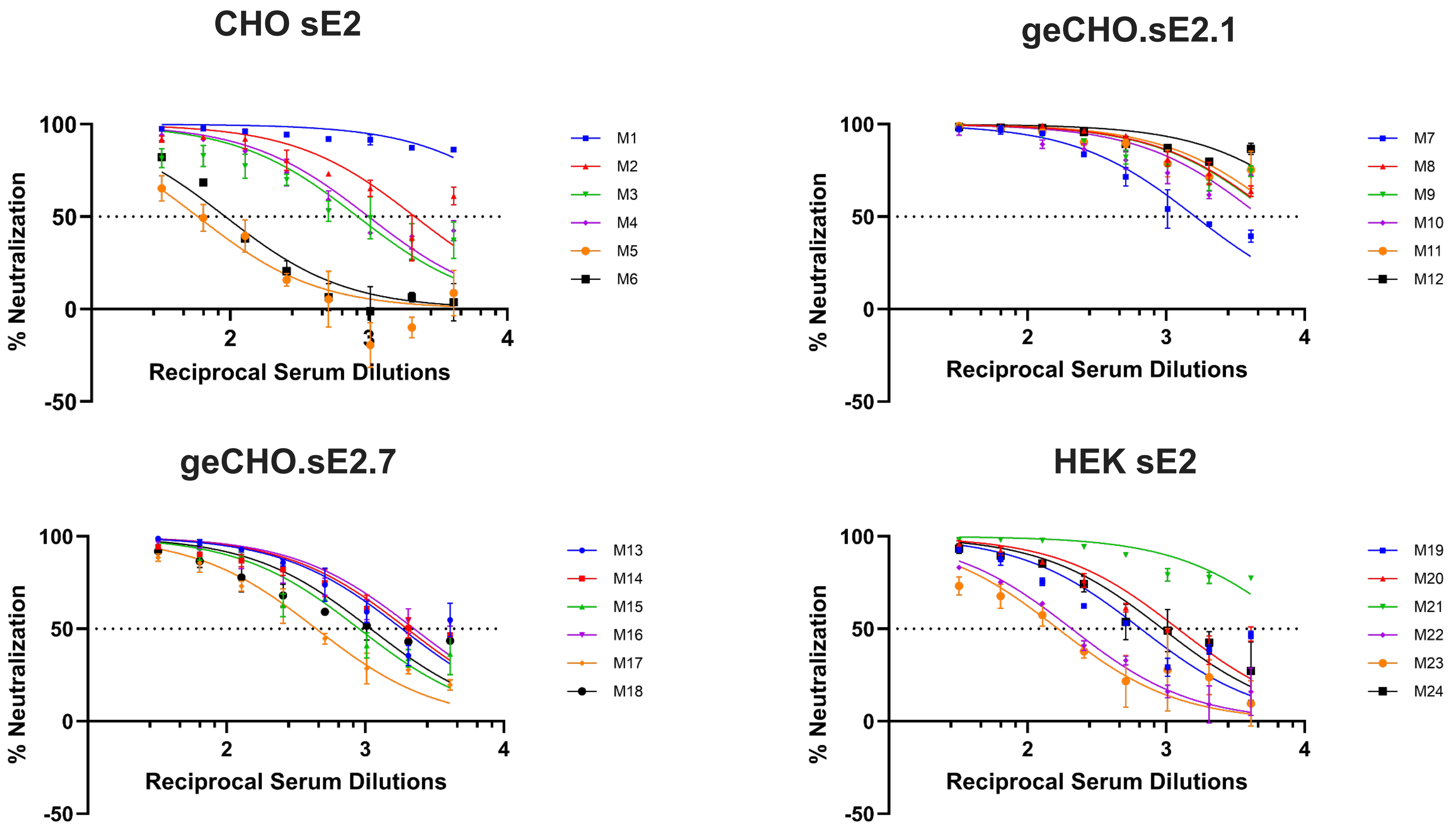


**Figure S7.** HCVpp neutralization curves depicting the percentage neutralization of the homologous isolate (H77C) for Day 56 serum dilutions of individual mouse sera for each group. The first dilution point was 1:32 and the sera were serially diluted twofold thereafter with the final dilution being 1:4096. The experiment was performed in duplicate and the error bars represent the standard error of the means (SEM).

### Table S1 - Antigenicity Analysis

| **Antibody** | **Domain** | **$K_d$ (nM)** | | | |
|---|---|---|---|---|---|
| | | CHO sE2 | geCHO.sE2.1 | geCHO.sE2.7 | HEK sE2 |
| CBH-4B | A | 0.85 ± 0.07 | 0.14 ± 0.01 | 0.30 ± 0.01 | 0.27 ± 0.01 |
| CBH-4D | A | 0.52 ± 0.05 | 0.11 ± 0.01 | 0.21 ± 0.01 | 0.18 ± 0.02 |
| AR3A | B | 1.7 ± 0.1 | **1.1 ± 0.2** | **1.0 ± 0.1** | 5.2 ± 0.1 |
| HEPC74 | B | 2.3 ± 0.5 | **1.4 ± 0.2** | **1.6 ± 0.1** | 8.0 ± 0.5 |
| HC84.1 | D | 0.26 ± 0.03 | **0.13 ± 0.01** | 0.37 ± 0.17 | 0.42 ± 0.14 |
| HC84.26.WH.5DL | D | 0.36 ± 0.07 | **0.14 ± 0.01** | 0.45 ± 0.04 | 0.56 ± 0.03 |
| HC33.1 | E | 3.6 ± 0.1 | 3.7 ± 0.1 | 3.7 ± 1.5 | 2.8 ± 0.1 |
| HCV1 | E | 2.4 ± 0.1 | 2.8 ± 0.1 | 2.5 ± 0.2 | 2.5 ± 0.8 |

### Table S2 – GT1a HCVpp $ID_{50}$ individual mice

| Mouse | Group | | | |
|---|---|---|---|---|
| | *CHO sE2* | *geCHO.sE2.1* | *geCHO.sE2.7* | *HEK sE2* |
| **M1** | 1,854 | 1,625 | 1,843 | 657 |
| **M2** | 2,145 | 6,268 | 2,008 | 1,219 |
| **M3** | 831 | 6,108 | 902 | 8,964 |
| **M4** | 996 | 4,846 | 2,257 | 208 |
| **M5** | 58 | 7,324 | 443 | 166 |
| **M6** | 92 | 14,374 | 1,096 | 944 |
| **Geometric mean** | 749 | 5626 | 1241 | 785 |
